# Mitochondrial Hyperactivity and Reactive Oxygen Species Drive Innate Immunity to the Yellow Fever Virus-17D Live-Attenuated Vaccine

**DOI:** 10.1101/2024.09.04.611167

**Authors:** Samantha G. Muccilli, Benjamin Schwarz, Forrest Jessop, Jeffrey G. Shannon, Eric Bohrnsen, Byron Shue, Seon-Hui Hong, Thomas Hsu, Alison W. Ashbrook, Joseph W. Guarnieri, Justin Lack, Douglas C. Wallace, Catharine M. Bosio, Margaret R. MacDonald, Charles M. Rice, Jonathan W. Yewdell, Sonja M. Best

**Affiliations:** Innate Immunity and Pathogenesis Section, Laboratory of Neurological Infections and Immunity, Rocky Mountain Laboratories, NIAID, NIH, Hamilton, M; Cellular Biology Section, Laboratory of Viral Diseases, NIAID, NIH, Bethesda, MD; Research Technologies Branch, NIAID, NIH, Hamilton, MT; Immunity to Pulmonary Pathogens Section, Laboratory of Bacteriology, NIAID, NIH, Hamilton, MT; Laboratory of Virology and Infectious Disease, The Rockefeller University, New York, NY; Center for Mitochondrial and Epigenomic Medicine, Children’s Hospital of Philadelphia, Philadelphia, PA; Integrated Data Sciences Section, Research Technologies Branch, NIAID, NIH

**Keywords:** Yellow fever virus, live-attenuated vaccine, mitochondria, electron transport, reactive oxygen species, type I interferon

## Abstract

The yellow fever virus 17D (YFV-17D) live attenuated vaccine is considered one of the successful vaccines ever generated associated with high antiviral immunity, yet the signaling mechanisms that drive the response in infected cells are not understood. Here, we provide a molecular understanding of how metabolic stress and innate immune responses are linked to drive type I IFN expression in response to YFV-17D infection. Comparison of YFV-17D replication with its parental virus, YFV-Asibi, and a related dengue virus revealed that IFN expression requires RIG-I-like Receptor signaling through MAVS, as expected. However, YFV-17D uniquely induces mitochondrial respiration and major metabolic perturbations, including hyperactivation of electron transport to fuel ATP synthase. Mitochondrial hyperactivity generates reactive oxygen species (mROS) and peroxynitrite, blocking of which abrogated IFN expression in non-immune cells without reducing YFV-17D replication. Scavenging ROS in YFV-17D-infected human dendritic cells increased cell viability yet globally prevented expression of IFN signaling pathways. Thus, adaptation of YFV-17D for high growth uniquely imparts mitochondrial hyperactivity generating mROS and peroxynitrite as the critical messengers that convert a blunted IFN response into maximal activation of innate immunity essential for vaccine effectiveness.

## Introduction

Yellow fever virus (YFV) is the prototypical *Orthoflavivirus*, positive-stranded RNA arboviruses that cause significant global morbidity and mortality each year [1]. Between the 15^th^ and 19^th^ centuries, yellow fever was among the most devastating infectious diseases in old and new worlds, including a 1793 outbreak that killed approximately 10% of Philadelphia’s population in the USA [2–4]. Walter Reed discovered YFV as the etiological agent early in the 20^th^ century. In rapid succession the *Aedes aegypti* mosquito was identified as its vector and the YFV-17D strain was generated as one of the most effective vaccines of all time [5–7]. Despite this, YFV remains an important human pathogen, causing approximately 80,000-200,000 infections annually with an estimated 40% mortality rate, mainly in Central and South America and Africa [8].

The YFV-17D vaccine was empirically derived by serial passage of a virulent YFV strain (YFV-Asibi) in mouse and chicken embryos [6, 7]. Administered to more than 500 million people worldwide, YFV-17D elicits strong innate and adaptive immune responses, and confers lifelong immunity to yellow fever in more than 95% of vaccinees [9]. YFV-17D robustly induces cytotoxic T cells, T_H_1 and T_H_2 CD4 T cells, and neutralizing antibodies that persist for 40 or more years [10, 11]. The strength and quality of the adaptive immune response is strongly influenced by detectable viremia [12] and a robust innate immune signature [13–15]. All three classes of interferons (IFNs) are linked to inducing and shaping the adaptive immune response to the virus, as well as controlling YFV-17D replication and dissemination *in vivo* [16–18]. While the vaccine is considered safe, a small number of vaccinees (1/250,000) develop yellow fever vaccine-associated viscerotropic disease or neurotropic disease [8, 19]. Patients with compromised type I IFN function are at a higher risk of developing disseminating infection [20, 21]. Therefore, the innate immune response is considered the cornerstone of the success of the YFV-17D vaccine in both control of virus replication and induction of adaptive immunity. The extent to which signaling pathways uniquely engaged by YFV-17D drive heightened IFN and inflammatory responses compared to the parental strain or other orthoflaviviruses is central to our study.

Several different pattern recognition receptors (PRRs) can be activated in response to infection with orthoflaviviruses to initiate an antiviral response. These include RIG-I-like receptors (RLRs), toll-like receptors (TLRs) and the DNA sensing pathway cGAS-STING. Previous work examining IFN responses to YFV-17D used mouse bone marrow-derived dendritic cells (DCs) and suggested that multiple TLRs are engaged to produce IFNs and other pro inflammatory cytokines [18]. However, YFV does not productively replicate in mouse cells including monocytes and DCs. This makes mice a poor model for understanding how innate responses are activated given that cytosolic RLRs are essential in initiating innate immune responses that control orthoflavivirus dissemination and tissue tropism, and license downstream adaptive immune responses in IFN-competent models [22].

RLRs signal through the adaptor protein mitochondrial antiviral signaling protein (MAVS) to induce expression of IFNs, chemokines and pro-inflammatory cytokines. MAVS is localized on mitochondria and peroxisomes, and its oligomerization on mitochondrial membranes is required for assembling the MAVS signalasome to activate IRF3, IRF7 and NFκB transcription factors [23]. Accordingly, viruses manipulate the structure and inter-organellar communication of the mitochondria, peroxisomes, and ER to dampen innate immune signaling and enhance replication. For example, the orthoflavivirus dengue virus (DENV) induces mitochondrial elongation and convoluted membrane formation resulting in reduced ER-mitochondria contact sites and dampened MAVS signaling [24]. Conversely, viral mitochondrial manipulation to meet the metabolic needs of viral replication and dampen MAVS-dependent signaling can result in the release of mitochondrial DNA sensed by cGAS-STING that activates the IFN response [25–27].

In addition to PRR signaling, metabolic dysregulation and cellular stress responses may contribute to immune responses to YFV-17D vaccination. In humans, symptomatic responses to YFV-17D vaccination correlated with ER stress, reduced tricarboxylic acid (TCA) cycle functionality, increased plasma levels of reactive oxygen species (ROS), and induction of cellular redox genes [28]. Furthermore, the magnitude of adaptive immune responses to vaccination can be predicted by gene expression involved in glycolysis and the integrated stress response [9, 29]. Together, these findings indicate a role for metabolic stress in activating the immune response to YFV-17D vaccination. However, mechanisms of metabolic stress and how they integrate with pathogen sensing and innate immunity in the context of YFV-17D replication are not known. Here, we provide a molecular understanding of the linkage between metabolic stress and innate immune responses to the YFV-17D vaccine that underpin its resounding success.

## Results

### Hepatotropic orthoflaviviruses induce distinct type I interferon dynamics independently of mitochondrial morphology

To understand the contribution of mitochondrial dynamics to YFV-17D activation of innate immune responses, we first compared multi-cycle virus growth kinetics and type I IFN expression between YFV-17D, YFV-Asibi, and DENV2 in HepG2 and Huh7 human hepatoma cell lines (Figure 1A, Supplemental Figure 1A). While the viruses attained similar peak titers in HepG2 cells, YFV-17D reached near peak titer in the first 24 hours, while YFV-Asibi and DENV2 required an additional 24 h. IFNβ was detected in YFV-17D-infected cell supernatants by 48 hpi, increasing for another day. Peak expression was at least 10-fold higher than present in YFV-Asibi or DENV2-infected cell supernatants, as previously reported [30]. Of note, Huh7 cells failed to secrete measurable IFNβ following YFV-17D or DENV2 infection despite being highly permissible to infection (Supplemental Figure 1A). Based on the robust IFN response, HepG2 cells were selected for further experiments. siRNA targeting critical adaptor proteins, MAVS and STING, was used to characterize the dominant pathway of IFNβ induction. Depletion of MAVS resulted in a near complete reduction of IFNβ secretion following infection with YFV-17D (Figure 1B) or DENV2 (Figure 1C). In contrast, reduced STING expression did not affect IFNβ following YFV-17D infection, but reduced IFNβ production late in DENV2 infection, consistent with existing literature that mitochondrial DNA release late in DENV infection is sensed by cGAS/STING to contribute to induction of type I IFN [31]. These results confirm RLR-dependent signaling is a dominant mechanism of orthoflavivirus sensing in non-immune cells.

**Figure 1:**
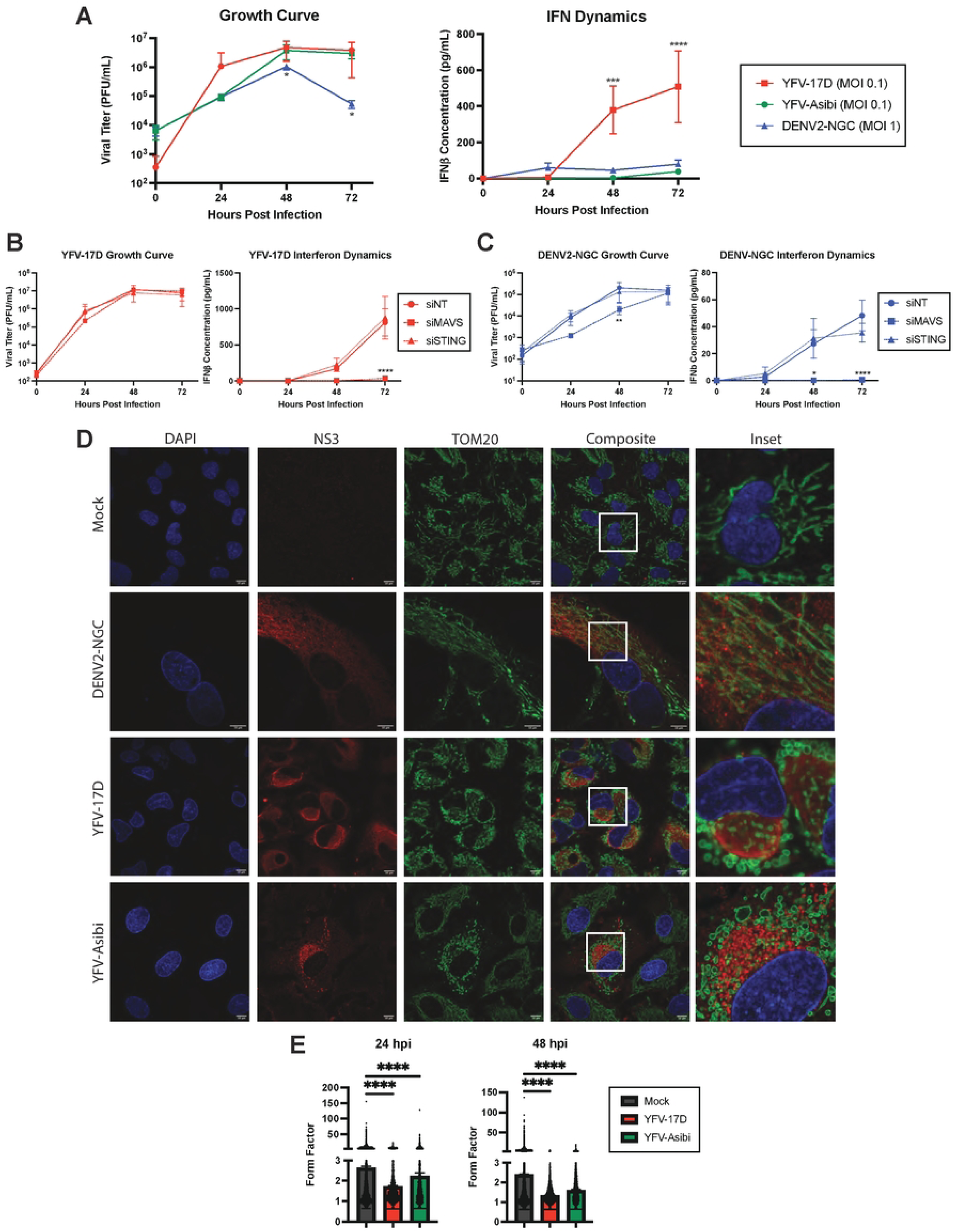
Dengue virus and yellow fever viruses modulate host mitochondrial and interferon dynamics differently. **A** Growth curve quantifying the viral titer and ELISA quantification of IFNβ secreted by HepG2 cells infected with YFV-17D (MOI 0.1), YFV-Asibi (MOI 0.1), or DENV2 (MOI 1) over a 72 hour period. **B and C** Growth curve quantifying the viral titer and ELISA quantification of IFNβ secreted by HepG2 cells infected with **B** YFV-17D or **C** DENV2 and treated with siRNAs to knockdown MAVS, STING, or a non-targeting control. **D** Immunofluorescence (IF) of mitochondrial morphology following infection with mock, DENV2 (MOI 1), YFV-17D (MOI 0.1), or YFV-Asibi (MOI 1) for 48 hours. Nuclei are stained with DAPI in blue, mitochondria are stained with TOM20 in green, and infected cells are stained with NS3 in red. **E** Quantification of the form factor of the mitochondrial network in HepG2 cells infected with mock, YFV-17D, or YFV-Asibi. Form factor was calculated using the following equation: Form Factor = (Perimeter)^2^ / 4π * Area. Values represent the mean ± SD (n=3). Statistical significance was assessed using two-way ANOVA followed by Tukey’s post hoc test for multiple groups (B and C) or ordinary one-way ANOVA followed by Tukey’s post hoc test for multiple groups (E). ****p<0.0001.

As orthoflaviviruses manipulate the structure and function of mitochondria, the predominant location of MAVS, we next examined the effects of viral infection on mitochondrial morphology via immunofluorescence staining for the outer membrane protein TOM20 and the viral nonstructural protein NS3 (Figure 1D). In mock-infected cells, mitochondria formed branched networks of intermediate length. As previously reported, DENV2 infection induced elongation and tangling of the mitochondrial network [24]. Conversely, infection with either YFV-17D or YFV-Asibi resulted in loss of branched mitochondrial networks in favor of individual, swollen mitochondria. Quantification of mitochondrial shape confirmed that YFV infection induced mitochondrial rounding at both 24hpi and 48hpi (Figure 1E). We extended these findings to Huh7 cells; DENV2 infection also induced mitochondrial elongation while YFV caused significant mitochondrial rounding and swelling (Supplemental Figure 1B).

Alterations in mitochondrial shape have a major impact on mitochondrial and cellular physiology. Mitochondrial elongation facilitates MAVS oligomerization and IFNβ production, while fission hinders MAVS’ ability to form high-order oligomers and impairs signaling [32, 33]. We next explored how manipulating mitochondrial networks modulates flavivirus induction of IFNβ using siRNA to deplete mitochondrial GTPases. Dynamin-related protein 1 (DRP1) has a major role in mitochondrial fission, while mitofusin 2 (MFN2) mediates mitochondrial fusion. As expected, DRP1 depletion elongated mitochondria while MFN2 depletion fragmented mitochondria in uninfected cells (Figure 2A). However, in YFV-17D infected cells fragmentation of mitochondria occurred under conditions of DRP1 depletion, suggesting that viral infection supersedes DRP1 dependence for mitochondrial fission. By contrast, mitochondrial hyperfusion in DENV2-infected cells was dependent on MFN2, as reported [24]. Virus replication was slightly increased (YFV-17D; Figure 2B) or was not significantly affected (DENV2; Figure 2C) by loss of MFN2 or DRP1. Surprisingly, knockdown of either protein resulted in significant loss of IFNβ production following infection with either virus (Figure 2B-2C). In contrast, in cells infected with Sendai virus (SeV, a paramyxovirus), a canonical inducer of MAVS-dependent IFNβ-expression, IFNβ secretion was increased in DRP1 siRNA-treated cells and decreased in cells depleted for MFN2 as has been previously published (Figure 2D). Together with the high IFNβ response by YFV-17D infected cells that display a fragmented mitochondrial phenotype, these data indicate that MAVS-dependent expression of IFNβ in response to infection with these hepatotropic flaviviruses is divorced from canonical mitochondrial morphology dynamics.

**Figure 2:**
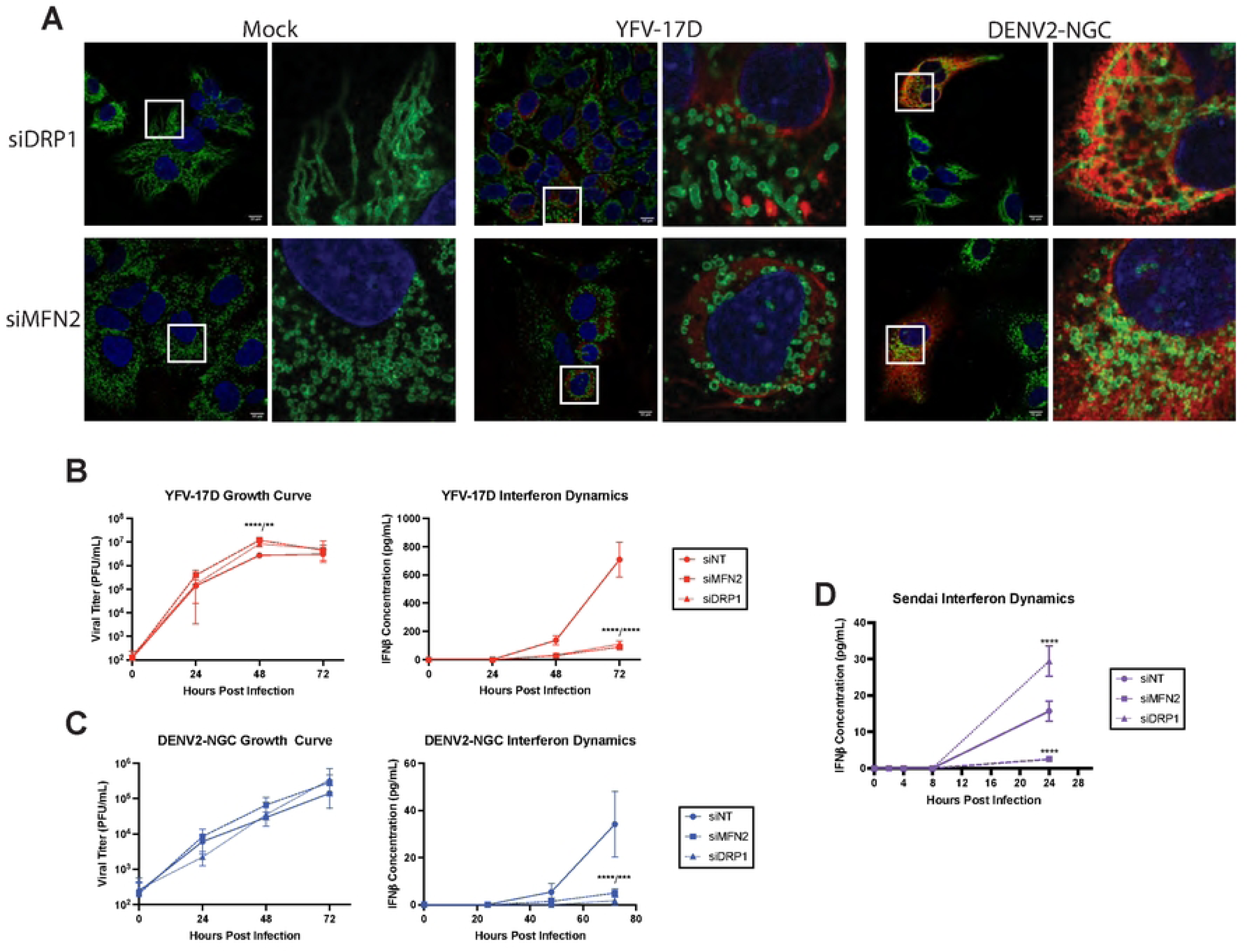
The type-I IFN response to YFV-17D and DENV2 is dissociated from mitochondrial morphology. **A** IF of mitochondrial morphology of HepG2 cells following infection with mock, YFV-17D (MOI 0.1), or DENV2 (MOI 1) and treated with siRNAs to knockdown DRP1 or MFN2. Nuclei are stained with DAPI in blue, mitochondria are stained with TOM20 in green, and infected cells are stained with NS3 in red. **B and C** Growth curve quantifying the viral titer and ELISA quantification of IFNβ secreted by HepG2 cells infected with **B** YFV-17D or **C** DENV2 and treated with siRNAs to knockdown MFN2, DRP1, or a non-targeting control. **D** ELISA quantification of IFNβ secreted by HepG2 cells infected with Sendai virus and treated with sirRNAs to knockdown MFN2, DRP1, or a non-targeting control. Values represent the mean ± SD (n=3). Statistical significance was assessed using two-way ANOVA followed by Tukey’s post hoc test for multiple groups (B-D). *** p<0.001 **** p<0.0001.

### YFV-17D drives rapid cellular energetic expenditure

We next used LC-MS-based metabolomics to determine how flavivirus-induced mitochondrial alterations correlate with metabolic changes in cells at 48 hpi. Principle component analysis (PCA) revealed that YFV-17D infection induced a distinct metabolic phenotype compared to mock, YFV-Asibi, or DENV2-infected cells (Figure 3A). Over 50% of measured metabolites were differentially abundant in YFV-17D infected cells compared to mock-infected cells with an FDR < 10% (Figure 3B). Conversely, YFV-Asibi and DENV2-infected cells had very few changes in metabolic intermediates relative to mock-infected cells despite major alterations in mitochondrial morphology (Figure 3A). Analysis of differentially abundant metabolites revealed that YFV-17D infection starkly depleted high-energy triphosphates in favor of the low energy monophosphates (Figure 3C, D), indicative of either energetic exhaustion or a high rate of energetic expenditure. Triphosphate pools were reduced to a lesser extent in YFV-Asibi or DENV2 infected cells. Metabolomic analysis performed 24 and 36 hpi revealed mock and YFV-17D infected cells were metabolically similar (Figure 3E). At these two timepoints, respectively, only two and five metabolites were differentially abundant between mock- and YFV-17D infected cells while triphosphate ratios were minimally different or unchanged (Figure 3F, Supplemental Figure 3). The tri/monophosphate ratios together with the lack of major metabolic changes indicate that this significant metabolic expenditure does not commence until after 36 hpi. Together, these data indicate that YFV-17D infection leads to rapid and intense metabolic expenditure that coincides with IFNβ secretion.

**Figure 3:**
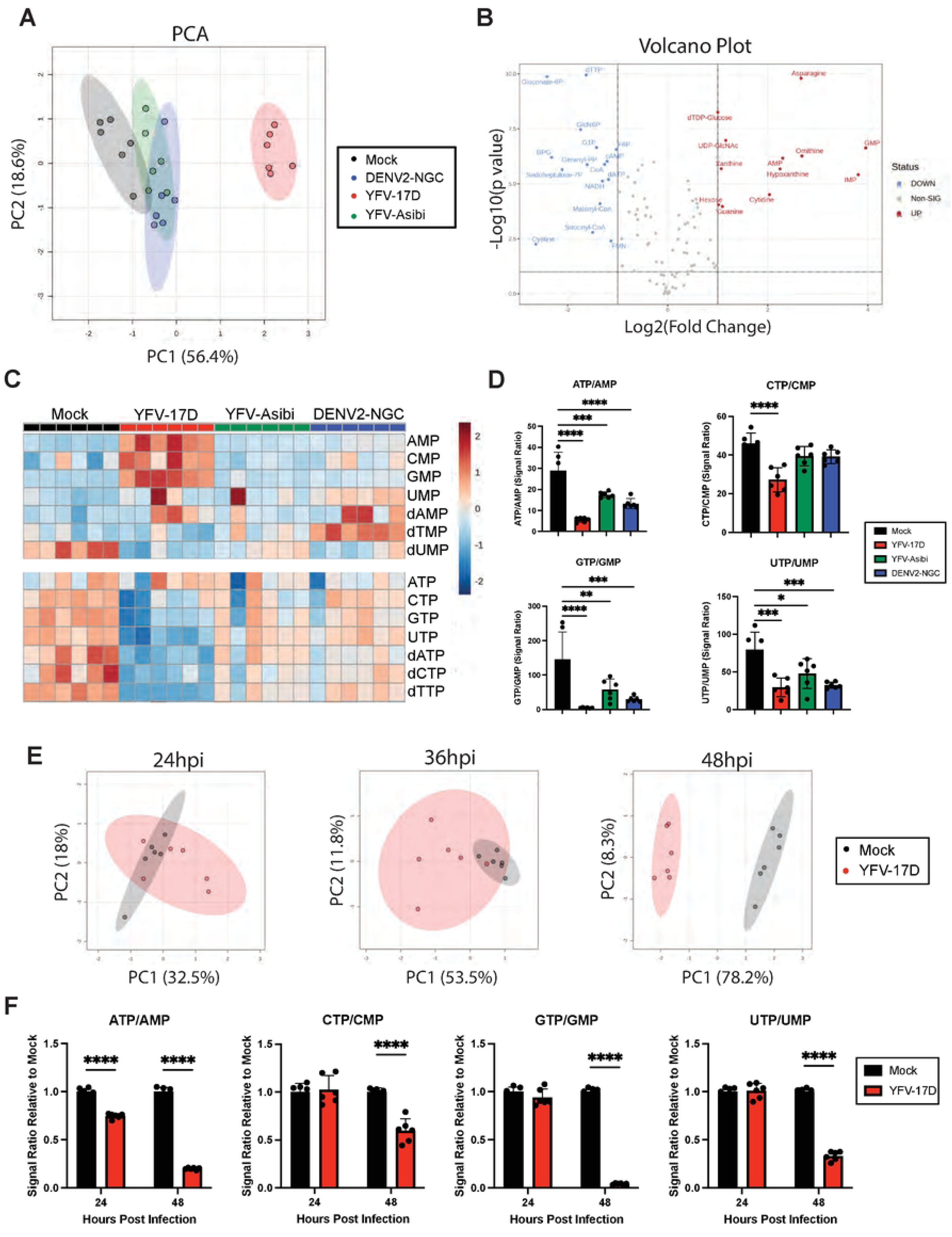
YFV-17D drives rapid cellular energetic expenditure. **A** Principle component analysis (PCA) of metabolomic differences between HepG2 cells infected with mock, DENV2 (MOI 1), YFV-17D (MOI 0.1), or YFV-Asibi (MOI 1) at 48hpi. **B** Volcano plots showing the significantly differentially abundant metabolites between YFV-17D infected HepG2 cells and mock-infected HepG2 cells at 48hpi as defined by metabolites with p<0.1 equivalent to an FDR of 10% via a Benjamini Hochberg. Metabolites with a fold change of ≥2 or ≤-2 are colored and named. **C** Metabolomic heatmap comparing the abundance of monophosphates and triphosphates following infection with mock, YFV-17D, YFV-Asibi, or DENV2. **D** Quantification of the triphosphate to monophosphate ratios in HepG2 cells infected with mock, YFV-17D, YFV-Asibi, and DENV2 at 48hpi. **E** PCA of metabolomic differences between mock and YFV-17D infected HepG2 cells at 24hpi, 36hpi, and 48hpi. **F** Quantification of the triphosphate to monophosphate ratios in mock and YFV-17D infected HepG2 cells at 24hpi and 48hpi. Values represent the mean ± SD (n=6). Statistical significance was assessed using ordinary one-way ANOVA followed by Dunnett’s multiple comparisons test (D) or two-way ANOVA followed by Sidak’s multiple comparisons test (F). **** p<0.0001.

### YFV-17D infection drives mitochondrial hyper-functionality and energetic overcommitment

We next used seahorse extracellular flux analysis to measure oxygen consumption over time in the presence of electron transport chain inhibitors to isolate the effects of infection on mitochondrial function. First, basal respiration is measured, then ATP-synthase is blocked by oligomycin to calculate the proportion of oxygen used for ATP synthesis. Next, the proton gradient is collapsed by FCCP that allows electrons to flow unimpeded across the inner mitochondrial membrane to measure maximal respiration rate. Spare respiratory capacity, a measurement of the cell’s ability to respond to an increase in energetic demands during metabolic stress, can be calculated by subtracting the maximal respiration rate from the basal respiration rate. Finally, a combination of complex I inhibitor rotenone and complex III inhibitor Antimycin A entirely abolish mitochondrial respiration, enabling calculation of non-mitochondrial respiration (Figure 4A-4B)[34].

**Figure 4:**
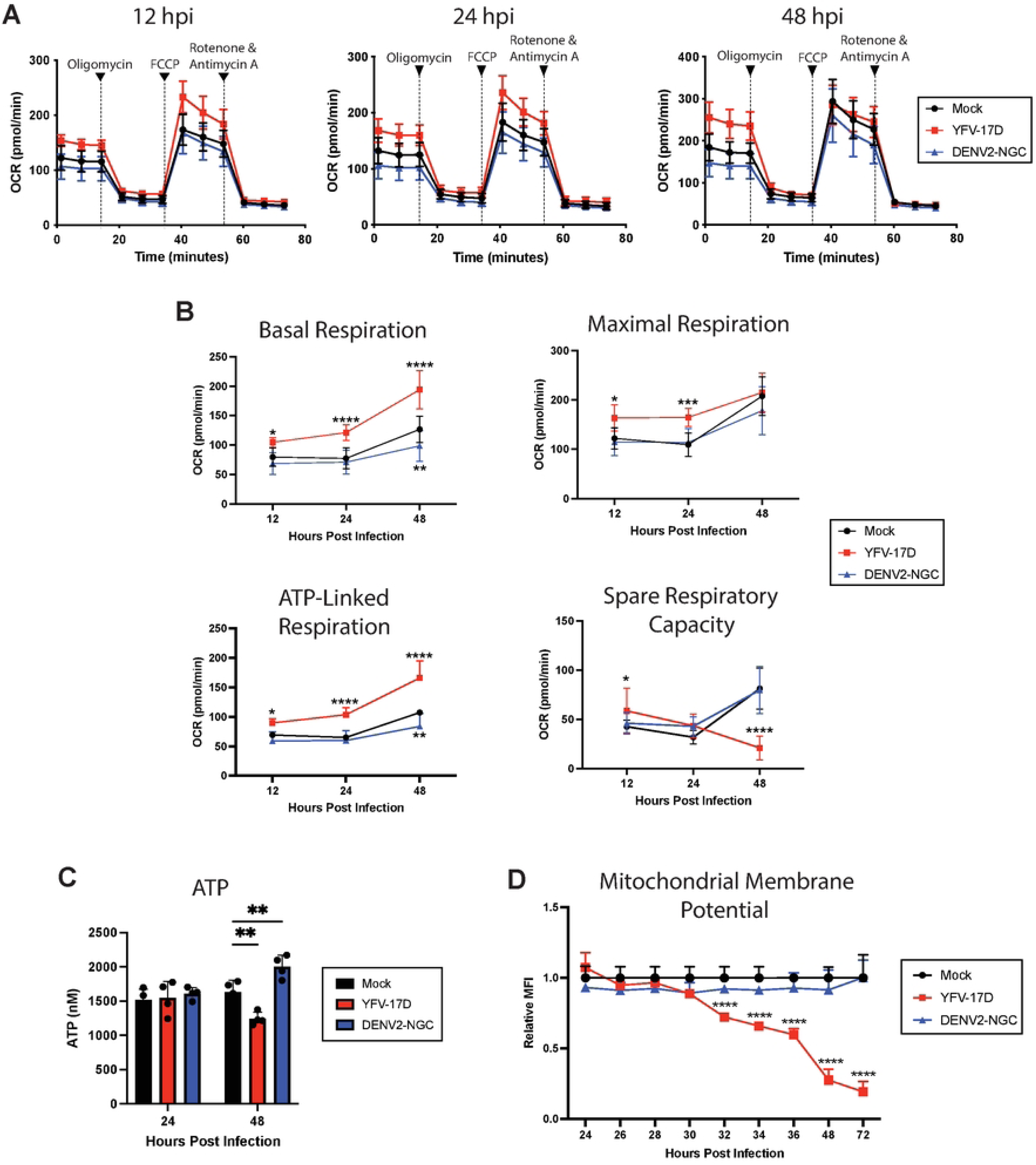
YFV-17D induces mitochondrial hyper-functionality and energetic overcommitment. **A** Seahorse XF cell mito stress test profile of HepG2 cells infected with mock, YFV-17D (MOI 0.1), or DENV2 (MOI 1) at 12hpi, 24hpi, and 48hpi. **B** Quantification of several Seahorse XF mito stress test parameters (basal respiration, maximal respiration, ATP-linked respiration, and spare respiratory capacity) in mock, YFV-17D, or DENV2 infected HepG2 cells over a 48 hour period. **C** Quantification of intracellular ATP levels in mock, YFV-17D, and DENV2 infected HepG2 cells at 24hpi and 48hpi. **D** Quantification of mitochondrial membrane potential (TMRE) in mock, YFV-17D, and DENV2 infected HepG2 cells between 24hpi and 72hpi. Values represent the mean ± SD (B: n=12, C: n=4, D: n=3). Statistical significance was assessed using two-way ANOVA followed by Dunnett’s multiple comparisons test (B, C, D). * p<0.05, ** p<0.01, *** p<0.001, **** p<0.0001.

Significantly increased basal respiration was evident as early as 12 hpi with YFV-17D. Notably, by 48hpi, the basal respiration rate of YFV-17D infected cells was nearly equal to the maximal respiration rate. ATP-linked respiration is the primary contributor to this basal respiratory elevation in the YFV-17D infected cells as represented by the difference between basal respiration and respiration following oligomycin addition. Further, YFV-17D infected cells increased the rate of glycolysis between 24 and 48 hpi as measured by metabolomics and Seahorse assays (Supplemental Figure 4A-C). Thus, YFV-17D infection maximizes mitochondrial respiration even under basal conditions, resulting in a significant decrease in mitochondrial spare respiratory capacity. In contrast, mitochondrial function was not changed by DENV2 infection, except for a decrease in basal respiration and glycolysis observed at 48 hpi.

In general, high mitochondrial respiration is associated with increased oxidative phosphorylation and ATP production. Yet, the metabolomics analysis indicated that YFV-17D infection depletes ATP. Therefore, we sought to verify the metabolomics by quantifying total ATP concentration in infected cells. Indeed, despite substantial increases in total- and ATP synthase-linked respiration at 48 hpi, YFV-17D infected cells had a significantly lower standing pool of ATP compared to DENV2- or mock-infected cells (Figure 4C). The concomitant increases in glycolysis and ATP synthase-linked respiration, drop in spare respiratory capacity, maintenance of maximal respiration, and drop in triphosphate pools suggest extremely high energetic demands in the YFV-17D infected cells that is outpacing the maximally available triphosphate synthesis.

While the seahorse analysis suggested increased mitochondrial function and health, the discordance of the flux data with ATP levels suggested that additional stressors and processes may be present in the mitochondria that were invisible to the measure of respiration under controlled conditions. To assess mitochondrial function in situ, we next measured mitochondrial membrane potential using flow cytometry. Mitochondrial membrane potential is driven by the electron transport chain pumping protons into the intermembrane space that are then used by ATP synthase to produce ATP. In a healthy cell, at equilibrium, the electron transport chain pumps more protons than ATP synthase consumes resulting in a high mitochondrial membrane potential. Interestingly, we found mitochondrial membrane potential decreased precipitously in YFV-17D infected cells beginning at 32hpi, while DENV2-infected cells maintained mitochondrial membrane potential (Figure 4D). This low mitochondrial membrane potential in conjunction with high ATP synthase-linked respiration indicates the mitochondrial depolarization is likely caused by ATP-synthase outpacing the ability of the electron transport chain to replenish the proton pool. This disconnect between electron transport and ATP synthesis is unsustainable and places the cell under significant metabolic stress. Together these data indicate that YFV-17D infection requires enormous amounts of ATP to keep up with the metabolic demands of infection. The cell makes every attempt to meet this extreme metabolic burden including upregulating both glycolysis and mitochondrial respiration. Thus, mitochondria become hyper-functional and energetically overcommitted, upregulating ATP synthase activity to the point that ATP synthesis surpasses the rate of mitochondrial respiration. While minimal cell death was apparent at the time of these measurements up to 48 hpi, the cell is eventually no longer able to maintain the rapid rate of mitochondrial respiration and ATP synthesis and succumbs to cell death by 72 hpi as measured by Annexin V staining (Supplemental Figure 5).

To alleviate concern that the major phenotypes that differentiate YFV-17D infection from its parental strain are a result of mutations acquired during tissue culture passage of the viruses, we repeated select assays using molecular viral clones. Recombinant YFV-17D and YFV-Asibi derived from molecular clones mirrored the tissue culture passaged lab stocks in growth kinetics, IFNβ secretion, and differential loss of mitochondrial membrane potential (Supplemental Figure 6A-C). Notably, although a dose-dependent effect of MOI was apparent for induction of IFNβ secretion and mitochondrial membrane potential loss during YFV-17D infection, increasing infection burden by YFV-Asibi did not increase IFNβ secretion or induce loss of mitochondrial membrane potential by 48 hpi (Supplemental Figure 6C). Thus, these phenotypes represent virus-encoded differences between YFV-17D and YFV-Asibi.

### YFV-17D-induced ROS is responsible for activating type I interferon signaling

In addition to the generation of ATP, mitochondrial respiration also generates mitochondrial reactive oxygen species (mROS) that can activate a pro-inflammatory state in cells [35, 36]. YFV-17D induced mROS over baseline levels measured by Mitosox labeling by 24 hpi whereas for DENV2, increased mROS required an additional 24 hours (Figure 5A-B). Treating cells with the cell permeable ROS scavenger MnTBAP at either 0 or 24 hpi did not impede replication of YFV-17D or DENV2. However, IFNβ production in YFV-17D infected cultures was reduced approximately 10-fold to levels equivalent to those produced following DENV2 infection (Figure 5C-D). MnTBAP treatment only delayed the induction of IFNβ by DENV2-infeccted cells. Importantly, this finding demonstrates that high levels of IFNβ secretion are separable from virus replication and accompanying high levels of viral molecular patterns and reveals that mROS is a critical factor required to drive high levels of IFNβ by YFV-17D.

**Figure 5:**
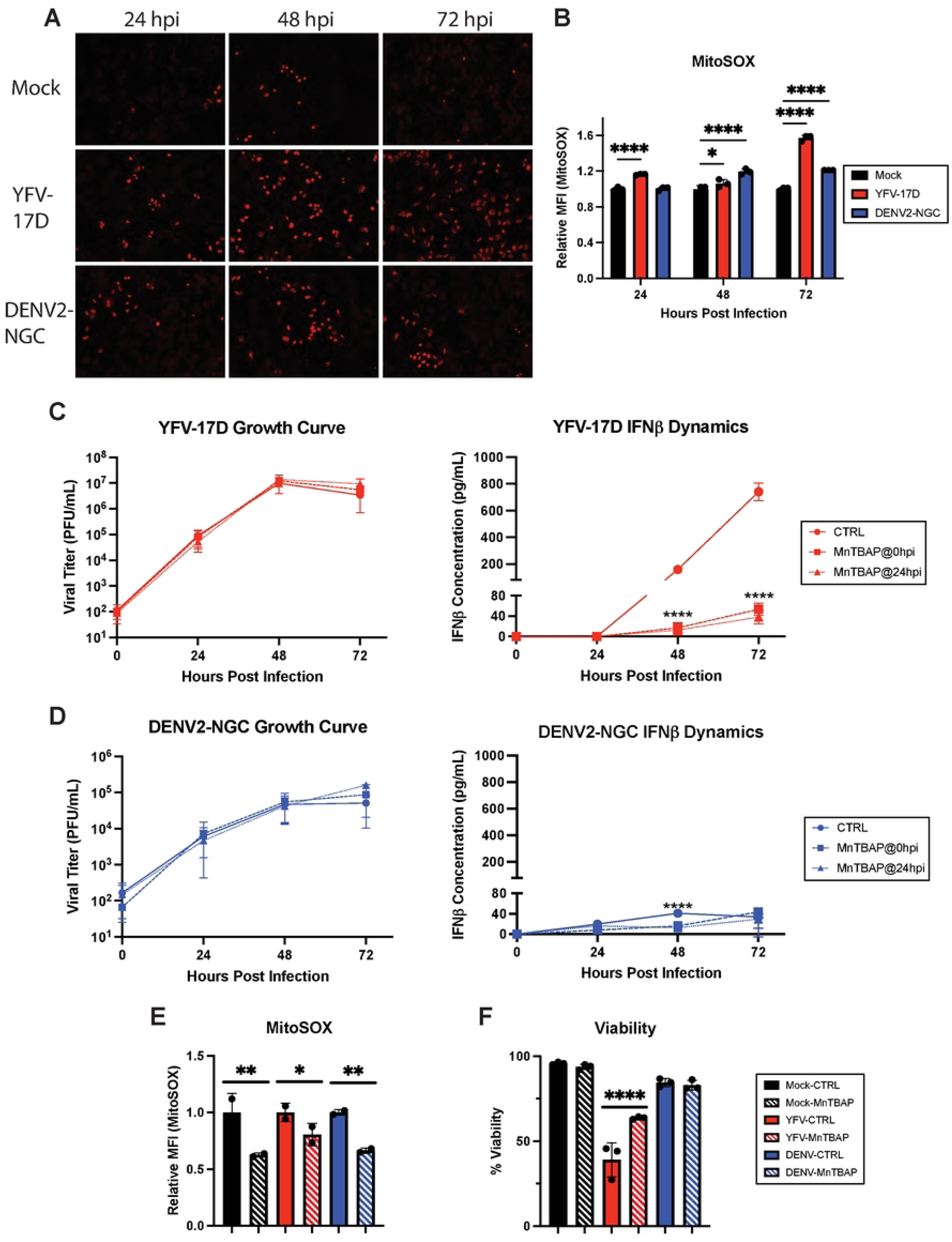
YFV-17D-induced ROS is responsible for activating the type-I IFN response and cell death during YFV-17D infection. **A** IF of mitochondrial ROS production using MitoSOX dye in HepG2 cells infected with mock, YFV-17D (MOI 0.1), or DENV2 (MOI 1) at 24hpi, 48hpi, and 72hpi. **B** Quantification of MitoSOX staining in mock, YFV-17D, and DENV2 infected HepG2 cells using flow cytometry. **C and D** Growth curve quantifying the viral titer and ELISA quantification of IFNβ secreted by HepG2 cells infected with **C** YFV-17D or **D** DENV2 and treated with MnTBAP [100µM] at 0hpi or 24hpi. **E** Flow cytometric quantification of MitoSOX staining in mock, YFV-17D, and DENV2 infected HepG2 cells treated with MnTBAP or a vehicle control for 48 hours. **F** Flow cytometric quantification of viability of mock, YFV-17D, and DENV2 infected HepG2 cells treated with MnTBAP or a vehicle control for 48 hours. Values represent the mean ± SD (B, C, D, F: n=3; E: n=2). Statistical significance was assessed using two-way ANOVA followed by Tukey’s post hoc test for multiple groups (B-D) or ordinary one-way ANOVA followed by Tukey’s post hoc test for multiple groups (E, F). * p<0.05, ** p<0.01, *** p<0.001, **** p<0.0001.

Given that the type I IFN response is incredibly energy intensive and that MAVS activation results in mitochondrial depolarization, we sought to determine if mitochondrial hyperactivity and energetic overcommitment was driven by a need to supply ATP for innate immune signaling or if this mitochondrial phenotype precedes IFN production. However, inhibition of IFNβ production using either MnTBAP or siRNA to deplete cells of MAVS did not rescue the loss of mitochondrial membrane potential following YFV-17D infection (Supplemental Figure 7). Thus, mitochondrial overcommitment is driven directly by YFV-17D replication rather than the cell-intrinsic, energy intensive process of type I IFN signaling.

MnTBAP also scavenges peroxynitrite, a ROS species generated from superoxide and nitric oxide [37]. Peroxynitrite induction closely mirrored that of IFNβ induction during YFV-17D replication (Supplemental Figure 8A), and MnTBAP significantly decreased cellular peroxynitrite (Supplemental Figure 8B). Inhibiting nitric oxide synthase with L-NAME (Supplemental Figure 8C) also significantly decreased IFNβ expression following infection, consistent with peroxynitrite being a major contributor to IFNβ secretion (Supplemental Figure 8E). Finally, since high levels of ROS are cytotoxic, we investigated the effect of MnTBAP on cell viability. Treatment with MnTBAP partially rescued YFV-17D infected cells from death, consistent with linkage of type I IFN induction and cell death via ROS generation (Figure 5F). Taken together, our data indicate that YFV-17D uniquely induces maximal mitochondrial respiration that is unable to keep up with the metabolic needs of the cell. This metabolic stress results in enhanced mROS and peroxynitrite generation which greatly amplifies MAVS-dependent IFN signaling while also mediating cell death.

### ROS is required for human dendritic cells to mount an innate antiviral response to YFV-17D infection

Dendritic cells (DCs) are professional antigen presenting cells that are both important targets of YFV infection and required for activation of naïve T cells. Given their critical role in vaccination of naïve individuals, we examined the role of ROS in virus-induced DC activation. Monocyte-derived DCs from five human donors were infected with YFV-17D in the presence of MnTBAP or vehicle control. While viral replication rates varied among donors (Figure 6A), IFNβ was detected in all infected cultures by 48 hpi (Figure 6B). As with HepG2 infection, MnTBAP treatment significantly decreased IFNβ expression and increased cell viability in all five donors without affecting viral replication (Figure 6C, A).

**Figure 6:**
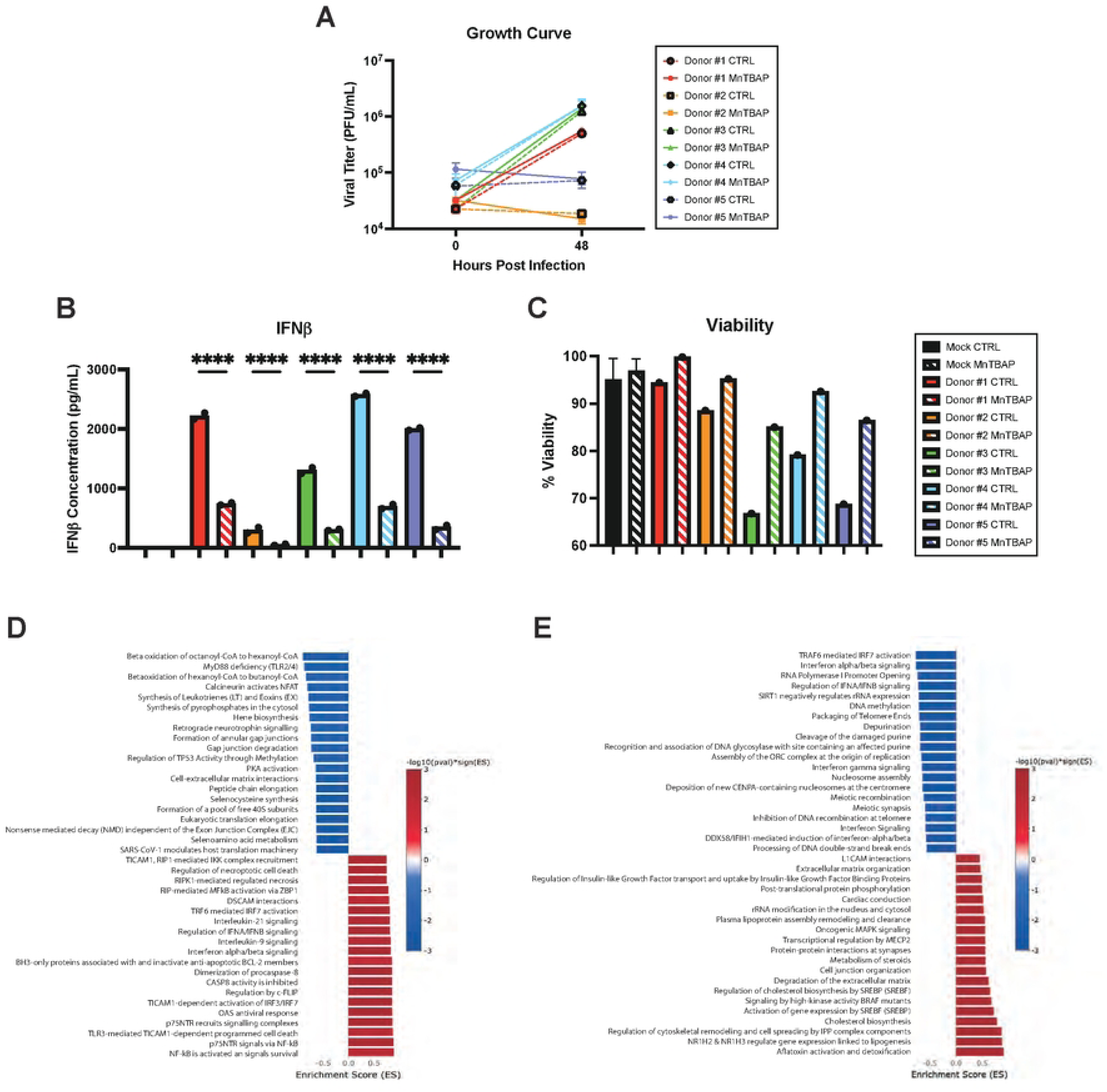
mROS is required for primary human dendritic cells to properly mount an innate immune response to YFV-17D infection. **A** Growth curve quantifying the viral titer of human DCs from five donors infected with YFV-17D (MOI 0.1) over a 48 hour period. **B** ELISA quantification of IFNβ secreted by DCs infected with either mock or YFV-17D. **C** Pathway enrichment analysis of YFV-17D infected DCs compared to mock infected DCs at 48hpi. **D** Pathway enrichment analysis of YFV-17D infected DCs treated with MnTBAP compared to vehicle control at 48hpi. Values represent the mean ± SD (n=2). Statistical significance was assessed using ordinary one-way ANOVA followed by Sidak’s multiple comparisons test (B). **** p<0.0001.

Global gene expression changes were assessed in DCs by bulk RNA sequencing at 48 hpi. Pathway enrichment analysis revealed that YFV-17D infection significantly upregulated mRNA involved in the innate immune response and cell death and downregulated protein translation associated mRNA (Figure 6D). MnTBAP treatment downregulated IFN signaling in infected cells (Figure 6E), with six of the top twenty downregulated pathways related to IFN signaling. Notably, multiple type I and III IFNs as well as interferon stimulated genes demonstrated ROS dependence (Supplemental Table 1). Global IFN signaling downregulation was accompanied by downregulating of genes involved in chemokine signaling (data not shown), pointing to a broad role for ROS signaling in chemotaxis and DC based T cell help.

Consistent with a central role for oxidative stress responses in cell-intrinsic responses to infection, key antioxidant enzymes (GSR, CAT, GPX1, and PRDX1) were downregulated by YFV-17D (Supplemental Figure 9A) and select genes (SOD2, NFE2L2, and KEAP1) were upregulated by MnTBAP treatment. Thus, YFV-17D infection likely increases oxidative stress by inducing mitochondrial ROS in concert with reducing cellular antioxidant capabilities. Together, our findings demonstrate that expression of type I and III IFNs and chemokines, considered hallmarks of YFV-17D vaccine responses, depend on a virus-induced metabolic program culminating in high ROS generation coincident with initiation of RLR signaling. ROS is therefore the critical secondary messenger that converts relatively low-level PRR signaling into the extraordinary induction of innate immunity following YFV-17D infection.

## Discussion

Compared to other licensed vaccines, including live-attenuated viruses and recombinant viral vectors, the YFV-17D vaccine induces unique immune signatures in human vaccinees associated with detectable viremia and delayed but robust innate responses [38]. The magnitude of neutralizing antibody and cytotoxic T cell responses correlate with the viremic load and antiviral cytokine and chemokine responses [12, 39]. In addition, cellular metabolic and stress responses enhance MHC I and II restricted antigen presentation and correlate with CD8 T cell responses [40, 41]. YFV-17D vaccine gene signatures have been predominantly ascribed to TLR stimulation [9, 18]. However, the dominant pattern recognition pathway controlling orthoflavivirus replication in IFN-competent animal models is MAVS-dependent [22]. Orthoflaviviruses also manipulate mitochondrial quality control pathways to facilitate replication and evade RLR signaling, amplifying the integrated stress and inflammatory responses [24, 42]. These fundamental features of orthoflavivirus infection and innate immune activation prompted us to reexamine the contribution of MAVS-dependent signaling and mitochondrial function to YFV-17D triggering of innate immunity as measured by IFNβ secretion.

As shown previously, YFV-17D replicates rapidly when compared to its parental strain YFV-Asibi or DENV2 [30]. Rapid replication is associated with major metabolic and mitochondrial perturbations. YFV-17D infection maximizes mitochondrial respiration to a rate that is energetically unsustainable. Indeed, our data indicate that YFV-17D infection causes ATP-synthase to consume protons faster than they can be supplied by the electron transport chain resulting in a rapid progressive reduction in mitochondrial membrane potential. Notably, this mitochondrial hyperactivity is independent of type I IFN signaling indicating that this is a process virally encoded rather than a side effect of an overactive innate immune response. Despite an extraordinary rate of ATP production as measured by highly elevated ATP-linked respiration, cells are still unable to meet the energetic demands of infection resulting in depletion of standing ATP pools. The onset of this metabolic stress is coincident with ROS generation and RLR activation. Thus, the metabolic stress of YFV-17D infection drives a reprioritization of nutrient/energetic resources to ROS generation and cytokine production prior to cell death. Importantly, blocking ROS-based signaling did not reduce viral replication. Thus even in the presence of high levels of viral molecular patterns, ROS is the key factor that translates relatively low RLR pathway activation into the YFV-17D hallmark innate immune signature. In human DCs, ROS signaling was critical for expression of type I and III IFNs, a spectrum of chemokines, ISGs, and cell death pathway activation, all which likely contribute to an adaptive immune response that confers lifelong protection from yellow fever. Importantly, this study reveals the cell biology of infection contributing to gene signatures associated with symptomatic responses and correlated with magnitude of adaptive immunity in humans following YFV-17D vaccination [9,13–15, 29].

Mitochondrial fusion, fission, and mitophagy are essential for maintaining cellular function. Viruses manipulate mitochondria to enhance replication through provision of metabolic intermediates for RNA replication as well as suppression of innate immune responses and apoptosis [43–45]. During RLR signaling, MAVS activation is aided by an intact mitochondrial membrane potential, and increased mitochondria surface area and ER-contacts [33, 44, 46]. Indeed, we confirmed important aspects of this model in the context of Sendai virus as a canonical RLR signaling activator. In Sendai virus infected cells, depleting MFN2 fragments mitochondria and reduces IFNβ secretion while depleting DRP1 elongates mitochondria and increases IFNβ secretion. However, while both YFV-Asibi and YFV-17D induced mitochondrial fragmentation, mitochondria in YFV-17D infected cells support high MAVS-dependent IFNβ secretion. We hypothesize that wild type YFV evolved to fragment mitochondria to dampen MAVS signaling to enhance replication and that this property was maintained in generating YFV-17D [47]. However, in the case of YFV-17D, infection-induced metabolic and oxidative stress negates any advantage provided by fragmented mitochondrial morphology. While YFV-induced mitochondrial fragmentation is independent of DRP1 (the major GTPase responsible for fission in uninfected cells), YFV-17D-induced IFNβ required both DRP1 and MFN2. MFN2-dependence is consistent with its role in adapting macrophage mitochondrial respiration to metabolic stress and promoting mROS generation [48]. The requirement for DRP1 in YFV-induced MAVS-dependent signaling is puzzling, particularly as DRP1 is inactivated by TBK1 to facilitate MAVS responses [49], necessitating further work to interpret this finding. Nevertheless, our results indicate that while mitochondrial morphology strongly contributes to MAVS-based anti-viral immunity, other factors can override its canonical role.

Viral-induced alterations in metabolism favoring glycolysis, as we observed in YFV-17D-infected HepG2 cells, likely evolve in part to increase nucleotide levels via the pentose phosphate pathway to support viral replication [50]. Similar metabolic reprogramming is critical in allowing virus-activated DCs to become optimal antigen-presenting cells. Increased glycolytic flux is required for TBK1-, IKKε- and Akt-dependent DC activation [51]. MAVS signaling is also regulated by glycolysis and activates the same kinases, consistent with these processes being linked in DCs [52]. Immune cell activation is also closely associated with ROS production. Similarly, mitochondrial hijacking by DENV, influenza A, and vesicular stomatitis virus results in ROS generation that enhances RLR and MAVS signaling [53, 54]. We confirmed a requirement for ROS in early expression of IFNβ by DENV-infected cells. While ROS is generally involved in RLR innate immune activation, the timing and magnitude of ROS-dependent IFNβ secretion points to critical differences between YFV-17D and related viruses.

Although the mechanism linking ROS to MAVS-dependent RLR signaling remains to be elucidated in the context of YFV-17D, several possibilities exist. Activated MAVS forms large, prion-like surface mitochondrial aggregates that are necessary and sufficient to activate downstream type I IFN signaling. It is well established that ROS-based MAVS aggregation generated by virus infection, chemicals, or systemic lupus erythematosus facilitates type I IFN signaling [55]. It is hypothesized that ROS oligomerizes MAVS by oxidizing specific cysteine residues, promoting disulfide bond formation and/or by peroxidizing lipids in the mitochondrial membrane, favoring MAVS transmembrane domain oligomerization [55, 56]. This might explain the critical role of mROS in YFV-17D induced IFNβ secretion, although there may be a contribution from ROS activation of NF-κB, AP-1, MAPK, and PI3K pathways [57]. Additionally, we show that YFV-17D infection induces the specific ROS species peroxynitrite, which via various alterations to proteins, lipids, sugars and nucleotides [58, 59], can independently influence metabolism and immune responses [60–62]. Additional studies are required to precisely define the contribution of ROS-based mechanisms to YFV-17D immune activation in liver cells and DCs.

Much also remains to be learned about how our findings relate to YFV-17D attenuation. Our data indicate that high metabolic burden is critical in augmenting the innate immune response to YFV-17D relative to YFV-Asibi and DENV2, which may be driven by rapid replication. YFV-17D differs from YFV-Asibi at 68 nucleotide positions, resulting in 32 non-synonymous codon changes scattered throughout the genome. Twelve changes are present in the viral envelope protein [63], conferring increased viral attachment and entry, increasing replication and enhancing proinflammatory pathway activation [30, 64]. This may contribute to our finding that robust YFV-17D replication leads to metabolic stress and mitochondrial overcommitment, culminating in ROS-induced type-I IFN signaling and cell death. In this case, paradoxically, high viral replication would be beneficial to the host as it leads to recognition and control of YFV-17D before significant viral dissemination can occur. Substitutions in non-structural proteins may also contribute to high RNA replication, or directly interfere with mitochondrial function. Indeed, mitochondrial manipulation has been attributed to nonstructural proteins NS4A and NS5 of DENV and ZIKV, respectively [24, 42]. Future reverse genetic-based mapping studies using the molecular clones used herein should elucidate the relevant mutations in YFV-17D responsible for its remarkable vaccine efficacy. This information will be useful for designing not only future flavivirus vaccines, but likely vaccines for a wide variety of viruses.

## Acknowledgements

This work was supported by the Division of Intramural Research, National Institutes of Health, National Institute of Allergy and Infectious Diseases, and R01AI124690 to CMR.

## Experimental Procedures

### Cell Culture

We cultured HepG2 cells (ATCC) and Vero E76 cells (ECACC) in DMEM (Gibco) supplemented with 10% FBS (Cytiva) and 1% penicillin/streptomycin (Gibco) at 37°C and 5% CO_2_. We cultured C636 mosquito cells (ATCC) in MEM (Gibco) supplemented with 10% heat-inactivated FBS, 2mM Glutamine (Gibco), 1% NEAA (Gibco), and 1% penicillin/streptomycin at 32°C and 5% CO_2_.

### Virus Infections

We used the following viruses in this study: yellow fever virus (YFV) strain 17D (from NIH Biodefense and Emerging Infections Research Recourses Repository, NIAID, NIH, NR116); YFV strain Asibi (from University of Texas Medical Branch World Reference Center for Emerging Viruses and Arboviruses); dengue virus (DENV) strain New Guinea C (from Dr. Adolfo García-Sastre); Sendai virus (SeV) strain Cantell (from Charles River Laboratories). We performed all procedures with YFV-Asibi under biosafety level-3 (BSL-3) conditions at the Rocky Mountain Laboratories Integrated Research Facility (Hamilton, MT). We propagated orthoflavivirus working stocks on C636 cells and titrated by plaque assay on Vero cells. Multiplicity of infection (MOI) is represented as plaque forming units (PFU) per cell. For all experiments, we infected cells for 1 hour at 37°C before removing virus inoculum and replacing with fresh culture medium.

### Immunofluorescence

We coated 4 well Lab-Tek II chamber coverglass slides (Thermo Fisher Scientific) with poly-l-lysine for 1 hour at 37°C prior to seeding cells. Following the culture period, we washed and fixed cells for 15 minutes at 37°C in 4% paraformaldehyde (Electron Microscopy Sciences) diluted in DMEM. Following fixation, we washed cells several times and permeabilized with 0.1% Triton 0/sodium citrate (Sigma Aldrich) for 15 minutes at room temperature before washing several more times. We blocked in 3% normal goat serum for 1 hour and then incubated overnight at 4°C in primary antibody diluted in 1% normal goat serum (Abcam). We washed cells and incubated with secondary antibody diluted in 1% normal goat serum for 1 hour at room temperature followed by Hoescht (Thermo Fisher Scientific) for 5 minutes at room temperature. We washed and stored cells at 4°C until imaging. We imaged processed slides using either a Zeiss LSM710 confocal microscope or a Leica Stellaris 8 confocal microscope and analyzed images using the FIJI software.

### Western Blot

We washed and lysed cells in SDS extraction buffer (50 mM Tris pH 7.4, 150 mM sodium chloride, 1mM EDTA, 2% SDS) supplemented with complete protease inhibitor (Sigma Aldrich). Lysates were agitated at 95°C for 10 minutes. We quantified protein concentration using the DC Protein assay (Bio-Rad) per the manufacturer’s protocol before diluting lysates 1:4 in sample buffer (90% 4X Protein Loading Buffer (LiCor), 10% DTT (Sigma Aldrich)). We heated diluted samples at 70°C for 10 minutes. We resolved 12 µg of cell lysate on 4-12% Bis-Tris gels (Thermo Fisher Scientific) and transferred to Nitrocellulose or PVDF membranes (Invitrogen) using the iBlot 2 Transfer System (Invitrogen). We blocked the membrane for 1 hour with Intercept Blocking Buffer (LiCor) followed by overnight incubation at 4°C in primary antibody diluted in blocking buffer. We washed membranes three times for 5 minutes with PBST before incubating at room temperature for 1 hour in secondary antibody diluted in blocking buffer. We washed membranes several times with PBST before imaging on the LiCor Odyssey CLx Imager.

### siRNA Treatment

We used the following siRNAs: Horizon ON-TARGETplus SMARTpool siRNAs specific against DRP1 (L-012092), MAVS (L-024237), MFN2 (L-012961), and STING (L-024333). We transfected HepG2 cells with 20 mmol siRNA using the Lipofectamine RNAiMAX reagent (Life Technologies) according to the manufacturer’s protocol. We incubated cells with siRNA proceed for 48 hours prior to infection and downstream analysis.

### Metabolite Sample Preparation

We washed HepG2 cells with a 0.9% sodium chloride solution at room temperature. To quench cellular metabolism, we added ice-cold LCMS-grade methanol (Fisher Chemical) and incubated on ice for 5 minutes. We added an equal volume of ice-cold LCMS-grade water (Fisher Chemical) before scraping cells and transferring samples to polypropylene tubes which were stored at -80°C until analysis. We randomized samples across plates to account for minor variations in sample collection time. The aqueous fraction was taken for LCMS analysis and diluted as needed in a 1:1 mixture of methanol and water to balance signal intensity.

### Liquid Chromatography Mass Spectrometry

We purchased tributylamine and all synthetic molecular references from Millipore Sigma. We purchased LCMS grade water, methanol, isopropanol and acetic acid through Fisher Scientific. We analyzed aqueous metabolites using a combination of two analytical methods with opposing ionization polarities [65, 66]. Both methodologies utilized a LD40 XR UHPLC (Shimadzu Co.) system for separation. Negative mode samples were separated on a Waters™ Atlantis T3 column (100Å, 3 µm, 3 mm X 100 mm) and eluted using a binary gradient from 5 mM tributylamine, 5 mM acetic acid in 2% isopropanol, 5% methanol, 93% water (v/v) to 100% isopropanol over 5 minutes. Signals were detected using a 5500 QTrap mass spectrometer (AB Sciex Pte. Ltd.). Two distinct MRM pairs in negative mode were used for each metabolite. Positive mode method samples were separated across a Phenomenex Kinetex F5 column (100 Å, 2.6 µm, 100 x 2.1 mm) and eluted with a gradient from 0.1% formic acid in water to 0.1% formic acid in acetonitrile over 5 minutes. Signals were detected using a 6500+ QTrap mass spectrometer (AB Sciex Pte. Ltd.). All signals were integrated using SciexOS 3.1 (AB Sciex Pte. Ltd.). Signals with greater than 50% missing values for a specific tissue set were discarded and remaining missing values were replaced with the lowest registered signal value. Where appropriate, signals with a QC coefficient of variance greater than 30% were discarded. Metabolites with multiple MRMs were quantified with the higher signal to noise MRM. The filtered dataset of the negative mode aqueous metabolites was total sum normalized after initial filtering. The positive mode aqueous metabolomics dataset was scaled and combined with the negative mode aqueous metabolite dataset using a common signal for serine. A Benjamini-Hochberg method for correction for multiple comparisons was imposed where indicated.

### Seahorse Extracellular Flux Analysis

We seeded Hep2G cells at 1 × 10^4^ cells per well in a XFe96 tissue culture plate (Agilent Technologies) and incubated for 24 hours in cDMEM. We infected cells with mock, YFV-17D (MOI 0.1) or DENV2 (MOI 1) as previously described. At 12-, 24-, and 48-hours post infection we performed extracellular flux analysis to interrogate the impacts of infection on mitochondrial function and glycolysis Prior to extracellular flux analyses, we washed cells twice with 200 μL of assay medium (minimal DMEM with 25 mM glucose, 2 mM sodium pyruvate, and 2 mM L-glutamine (Agilent Technologies)). We incubated cells in 180 µL Assay medium for 1 hour at 37°C in a non-CO_2_ incubator. We measured oxygen consumption rate (OCR) rate at baseline and following injection of oligomycin A1 (2 μM, MilliporeSigma), fluoro-carbonyl cyanide phenylhydrazone (FCCP; 2 μM; Cayman Chemical), and rotenone/antimycin (0.5 μM final concentration for both; MilliporeSigma). We used corresponding extracellular acidification rates (ECAR) as surrogate markers of changes in glycolytic flux. We performed all extracellular flux assays on the Seahorse XFe96 Analyzer (Agilent Technologies).

### Flow Cytometry

We used the following cellular dyes: TMRE (Abcam), MitoSOX (Abcam), Peroxynitrite Green (Abcam), and Annexin V/PI (Thermo Fisher Scientific) according to the respective manufacturer’s protocol. Following staining, we washed cells several times with PBS and split into single cell solution using trypsin. When appropriate, we also stained cells with the Live/Dead Fixable Aqua Dead Cell Stain (Thermo Fisher Scientific) for 20 minutes on ice to identify the live cell population. Flow cytometry analysis was performed on a BD LSR Fortessa X-20 flow cytometer.

### ATP Quantitation

We extracted ATP from cells using the boiling water method to minimize ATPase depletion. We washed HepG2 cells twice with PBS and then added boiling highly purified water (ThermoFisher), triturating to lyse cells and boiling for several more minutes to completely inactivate ATPases. We stored samples at -80°C until analysis. We measured in ATP in each sample in triplicate using an ATP Determination Kit (Thermo Fisher Scientific) according to the manufacturer’s protocol, analyzing all samples in one assay to maximize reproducibility.

### Small Molecule Inhibitors

We used MnTBAP (Sigma Aldrich) and L-NAME (Selleck Chemicals). We prepared a 50 mM stock solution of MnTBAP in DMSO (ATCC). Immediately before use, we prepared a reduced working solution by diluting the MnTBAP to a final concentration of 1mM in PBS/4% NaOH [2.5M] (Sigma Aldrich). We added 100µM of the reduced working solution to cells in a drop-wise fashion to minimize cell death. For this treatment, the vehicle control consisted of an equal volume of DMSO in PBS/4% NaOH [2.5M]. Alternately, we treated cells with 5mM L-NAME in sterile water and an equal volume of water was used as a vehicle control for these experiments. For both inhibitors, we retreated cells every 24 hours to maintain high inhibitor concentration.

### Full-length molecular clones

Full length YFV Asibi virulent strain (MT093734.1) and 17D vaccine strain (MT114401.1) infectious clones have been previously described [67, 68] and are designated pACNR-2015FLYF-Asibi and pACNR-2015FLYF-17Da respectively. The E.coli strain MC1061 was used to amplify these clones. After amplification and purification, plasmid DNA was linearized using AflII, followed by clean up and in vitro transcription using the mMESSAGE mMACHINE SP6 transcription kit. The virus stock was produced in Huh7.5 cells using 10μg of RNA essentially as described [69]. Cells of the electroporation reaction were seeded in a T-175 flask. At 4-6 hours post-electroporation, the media was changed and at 20-24 hours post-electroporation, media was replaced with serum free DMEM. Virus stocks were obtained by harvesting the media after an additional 6-8 hours and aliquots were stored at -80°C.

### Human monocyte-derived dendritic cells

Human monocyte-derived dendritic cells were prepared as previously described [70]. Briefly, we isolated peripheral blood mononuclear cells from human buffy coats (BioIVT) by centrifugation through a Ficoll-Paque Plus density gradient (Cytiva). Cells were enriched for CD14^+^ monocytes using a RossetteSep Monocyte Enrichment Kit (StemCell Technologies). Monocytes were resuspended at 1×10^6^ cells/mL in DC medium [RPMI + Glutamax (Invitrogen), 5% FBS, 15 mM Hepes, 0.1 mM nonessential amino acids, 1 mM sodium pyruvate, 100 units/mL penicillin and 100 μg/mL streptomycin] containing IL-4 (20 ng/mL) and GM-CSF (20 ng/ml) (PeproTech) and cultured for 5d with replacement of half of the medium and addition of fresh cytokines every other day. Nonadherent DC were harvested by centrifugation and suspended in DC medium. The resulting cells were determined to be >95% CD11c^+^/CD209^+^ by flow cytometry.

### Bulk RNA-seq of human DCs

We plated human monocyte-derived DCs at 2×10^6^ cells/well and mock-infected or infected with YFV-17D at an MOI of 0.1 in a small volume for 1 hour at 37°C, 5% CO_2_ with occasional gentle rocking. We added fresh DC medium to the wells to the desired volume and MnTBAP (50 µM) or vehicle control was added to the appropriate wells at 0 and 24 h post-infection. At 48 hpi, we harvested hDCs by centrifugation, and lysed the cells with 1 mL of TRIzol reagent/sample (Thermo Fisher Scientific), and samples were stored at -80°C before extraction and bulk RNA-seq. We collected cell supernatants at 0 and 48 hpi for cytokine and viral plaque assays.

For RNA extraction, we mixed the samples with 200 µL of 1-bromo-3-chloropropane (Sigma Aldrich) and centrifuged at 4°C for 15 minutes at 16,000 x g. We removed the aqueous phase and passed it through a QIAshredder column (Qiagen) at 21,000 x g for 2 minutes. RNA was extracted using the Qiagen AllPrep DNA/RNA system. An additional on-column DNase I treatment was performed for all extractions. We assessed RNA quality using the Agilent 2100 Bioanalyzer RNA 6000 Pico kit (Agilent Technologies) and quantified the RNA using the Quant-it RiboGreen RNA assay (Thermo Fisher Scientific).

Following RNA extraction, two hundred nanograms of RNA was used as the input for the Illumina stranded mRNA prep, ligation kit (Illumina) to generate sequencing libraries following the manufacturer’s protocol. The final libraries were analyzed using the Agilent bioanalyzer and the libraries were quantified using the Kapa SYBR FAST Universal qPCR kit for Illumina sequencing (Kapa Biosystems). The individual libraries were diluted to 6 nM and 2 uL of each added to a library pool. The library pool was denatured and diluted to a 10 pM stock and paired-end 2 x 75 cycle sequencing carried out on the MiSeq using a Nano V2 flow cell and 300 cycle chemistry. Following the sequencing run, reads per microliter mapping to human genes were determined for each library. The library pool was rebalanced and quantified using the Kapa SYBR FAST Universal qPCR kit for Illumina sequencing. The library pool was diluted to 9 pM and paired-end 2 x 75 cycle sequencing carried out on the MiSeq using a Nano V2 flow cell and 300 cycle chemistry. The final library pool containing 6 nM of each library was sent to the National Cancer Institute, Center for Cancer Research Sequencing Facility (NCI CCR-SF) for further sequencing. The samples were paired-end 2 x 100 cycle sequenced using a NovaSeq 6000 instrument and SP flow cell and 300 cycle chemistry.

Following sequencing, Raw fastq files were trimmed to remove adapters and low-quality bases using Cutadapt v1.18 before alignment to the GRCh38 reference genome and the Gencode v42 genome annotation using STAR v2.7.9a [71]. PCR duplicates were marked using the MarkDuplicates tool from the Picard v3.1.0 (https://broadinstitute.github.io/picard/) software suite. Raw gene counts were generated using RSEM v1.3.3 [72] and were filtered to include genes with >=1 count per million (CPM) in at least 2 samples.

Differential expression was evaluated using limma with TMM normalization [73]. For all differential expression comparisons, pre-ranked geneset enrichment analysis (GSEA) [74] was performed using the Reactome [75] and MitoCarta v3.0 [76].

### Statistical Analysis

All graphical values are shown as mean ± standard deviation. We performed statistical analyses using GraphPad Prism 10. Ordinary one-way or two-way ANOVAs with Dunnett’s multiple comparisons tests were performed, when appropriate, to test statistical significance between means. We considered p-values less that 0.05 to be statistically significant.

## Figure Legends

**Supplemental Figure 1:**
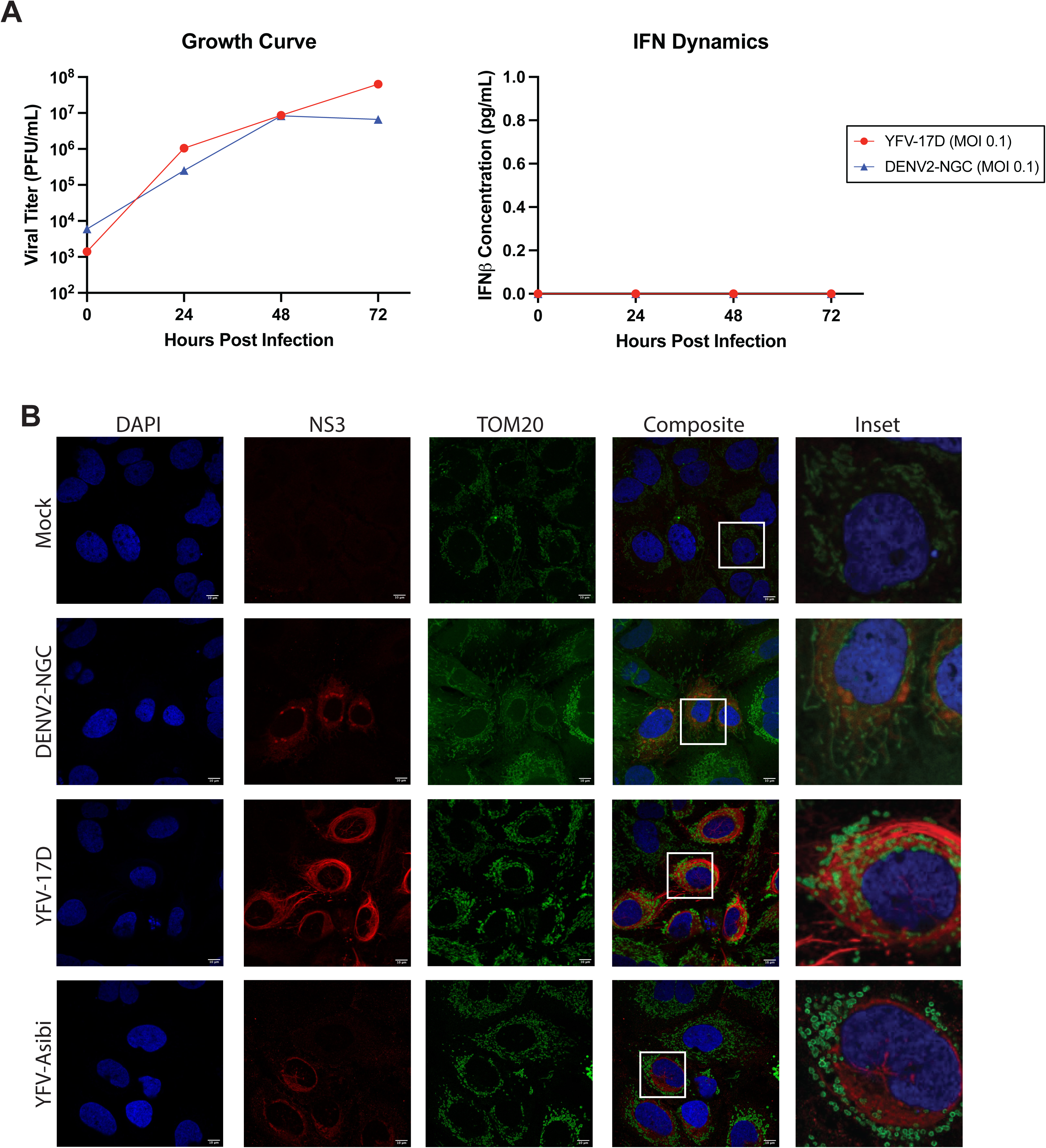
Differential mitochondrial morphodynamics are consistent across multiple cell types. **A** Growth curve quantifying the viral titer and ELISA quantification of IFNβ secreted by Huh7 cells infected with YFV-17D (MOI 0.1) or DENV2 (MOI 0.1) over a 72 hour period. **B** IF of mitochondrial morphology in Huh7 cells following infection with mock, DENV2 (MOI 1), YFV-17D (MOI 0.1), or YFV-Asibi (MOI 1) for 48 hours. Nuclei are stained with DAPI in blue, mitochondria are stained with TOM20 in green, and infected cells are stained with NS3 in red.

**Supplemental Figure 2:**
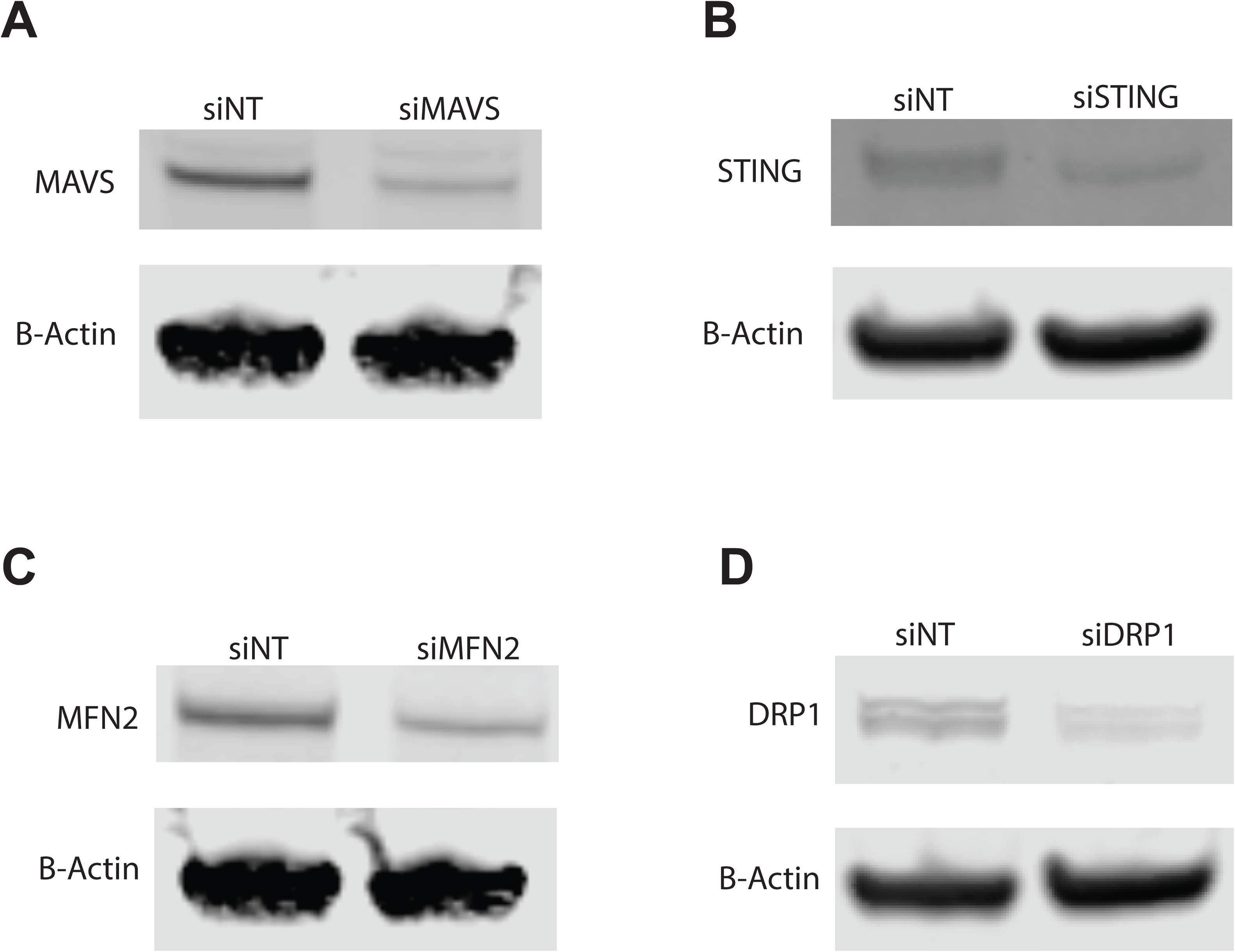
siRNA treatment results in knockdown of host. *proteins* Western blot quantifying protein following siRNA treatment with **A** siMAVS, **B** siSTING, **C** siMFN2, and **D** siDRP1. Treatment with siRNA resulted in significant knockdown of the respective protein.

**Supplemental Figure 3:**
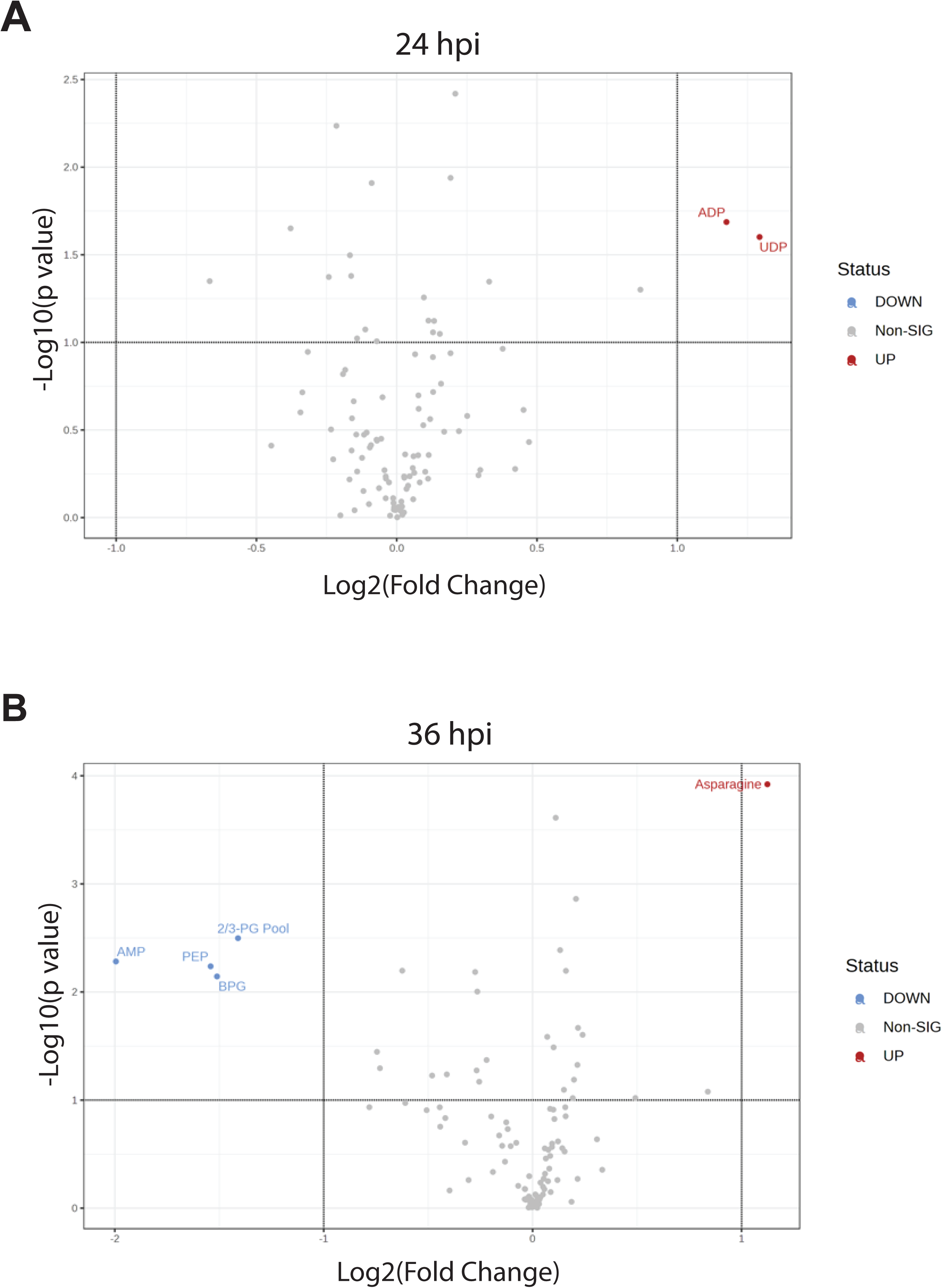
Very few metabolites are differentially abundant between YFV-17D and mock infected cells at 24 and 36 hpi. **A and B** Volcano plots showing the significantly differentially abundant metabolites between YFV-17D infected Hep G2 cells and mock-infected HepG2 cells at **A** 24hpi and **B** 36hpi.

**Supplemental Figure 4:**
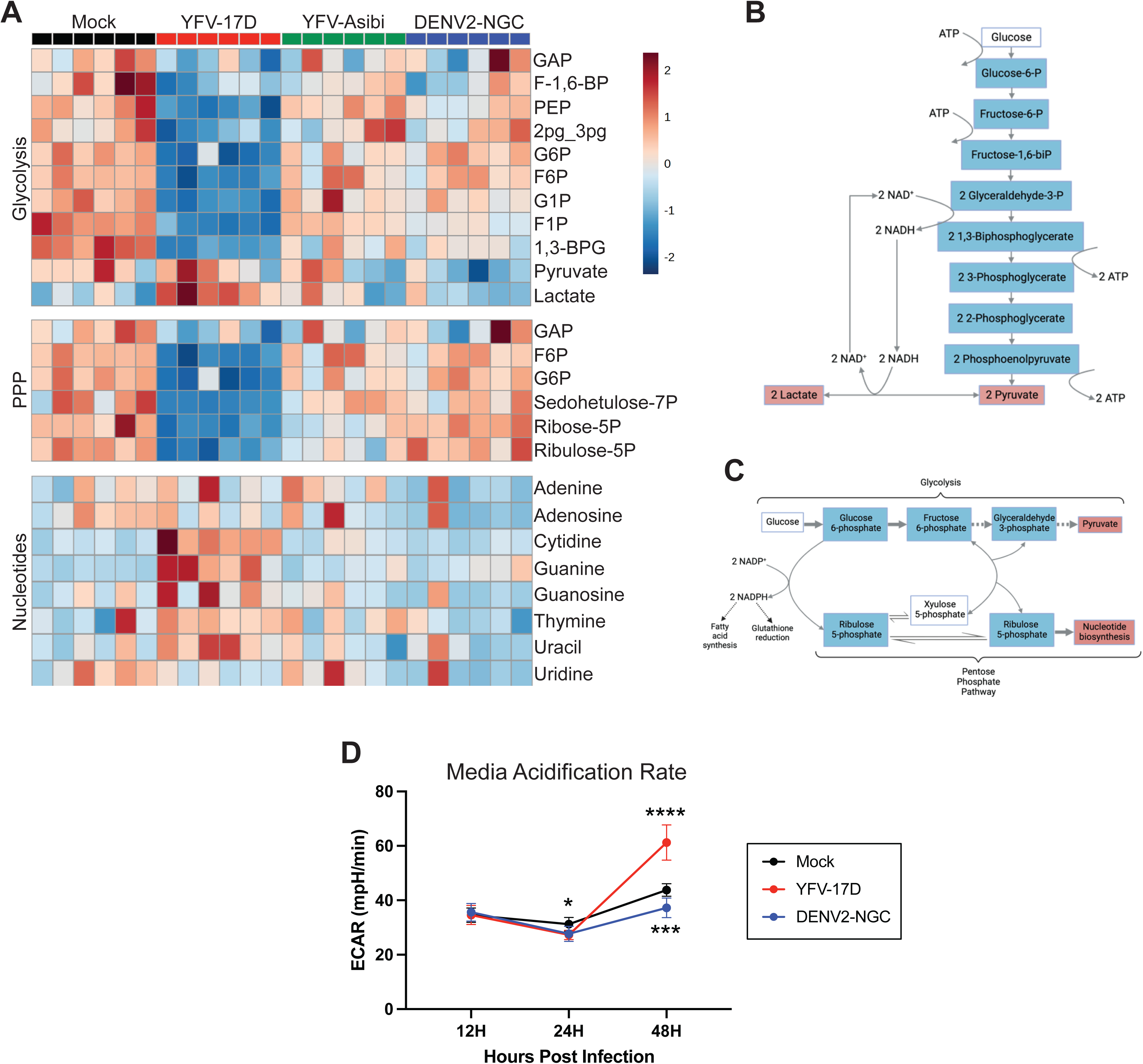
YFV-17D upregulates glycolysis between 24 and 48 hpi. **A** Metabolomic heatmap comparing the abundance of metabolites from the glycolytic pathway, pentose-5-phosphate pathway, and nucleotides following infection with mock, YFV-17D (MOI 0,1), YFV-Asibi (MOI 1), or DENV2 (MOI 1) in HepG2 cells. **B** Schematic of the glycolysis pathway with metabolites significantly downregulated in YFV-17D infection at 48hpi shown in blue and metabolites significantly upregulated in YFV-17D infection at 48hpi shown in red. **C** Schematic of the glycolysis and pentose-5-phosphate pathway with metabolites significantly downregulated in YFV-17D infection at 48hpi shown in in blue and metabolites significantly upregulated in YFV-17D infection at 48hpi shown in red. **D** Quantification of the media acidification rate obtained during the Seahorse XF mito stress test in mock, YFV-17D, or DENV2 infected HepG2 cells over a 48 hour period. Values represent the mean ± SD (n=12). Statistical significance was assessed using two-way ANOVA followed by Dunnett’s multiple comparisons test (D). * p<0.05, ** p<0.01, *** p<0.001, **** p<0.0001.

**Supplemental Figure 5:**
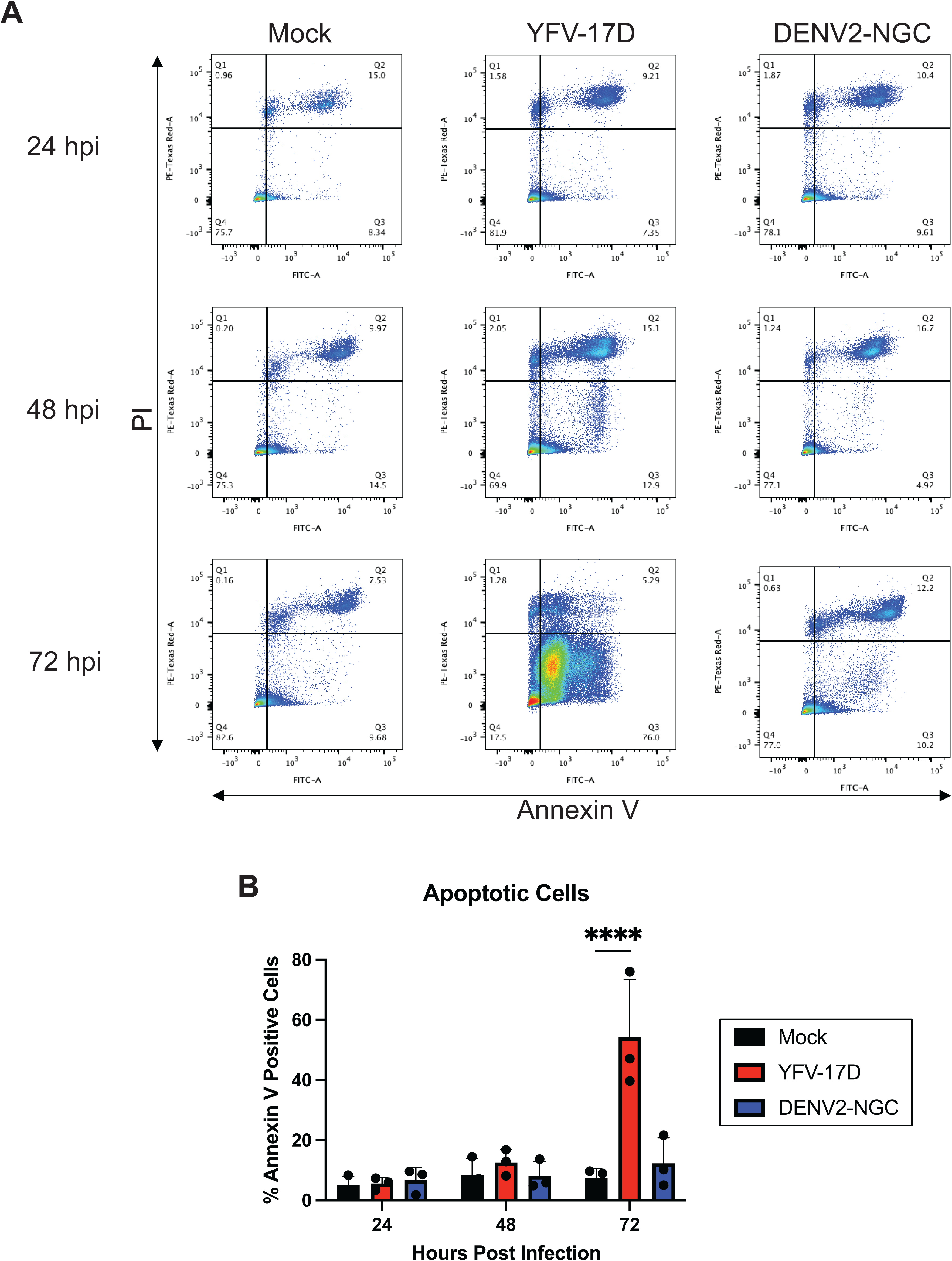
YFV-17D does not induce apoptosis until late timepoints. **A** Representative flow cytometry dot plots quantifying Annexin V and propidium iodide in mock, YFV-17D (MOI 0.1), and DENV2 (MOI 1) infected HepG2 cells at 24hpi, 48hpi, and 72hpi. **B** Quantification of the percentage of apoptotic cells in mock, YFV-17D, and DENV2 infected cells at 24hpi, 48hpi, and 72hpi. Values represent the mean ± SD (n=3). Statistical significance was assessed using two-way ANOVA followed by Dunnett’s multiple comparisons test (D). **** p<0.0001.

**Supplemental Figure 6:**
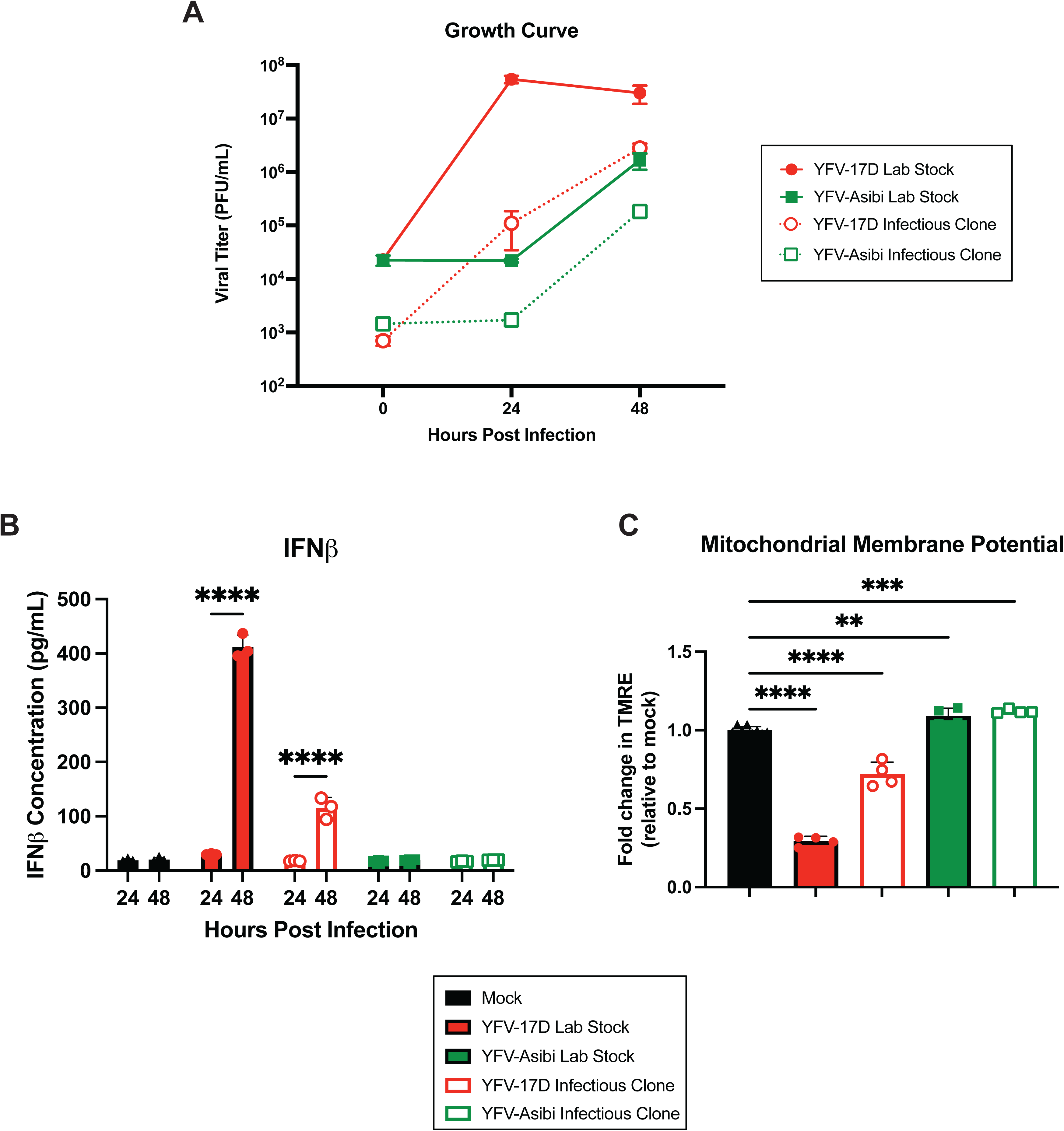
Infectious clones mirror the phenotypes of lab stock viruses. **A** Growth curve quantifying viral titer of HepG2 cells infected with YFV-17D (MOI 0.1) or YFV-Asibi (MOI 0.1) generated from lab stocks or infectious clones over a 48 hour period. **B** ELISA quantification of IFNβ secreted by HepG2 cells infected with lab stocks or infectious clones at 24 and 48 hpi. **C** Quantification of mitochondrial membrane potential (TMRE) in HepG2 cells infected with lab stock or infectious clone viruses at 48hpi. Values represent the mean ± SD (A, B: n=3; C: n=4). Statistical significance was assessed using two-way ANOVA followed by Dunnett’s post hoc test for multiple groups (B) or ordinary one-way ANOVA followed by Sidak’s post hoc test for multiple groups (C). ** p<0.01, *** p<0.001, **** p<0.0001.

**Supplemental Figure 7:**
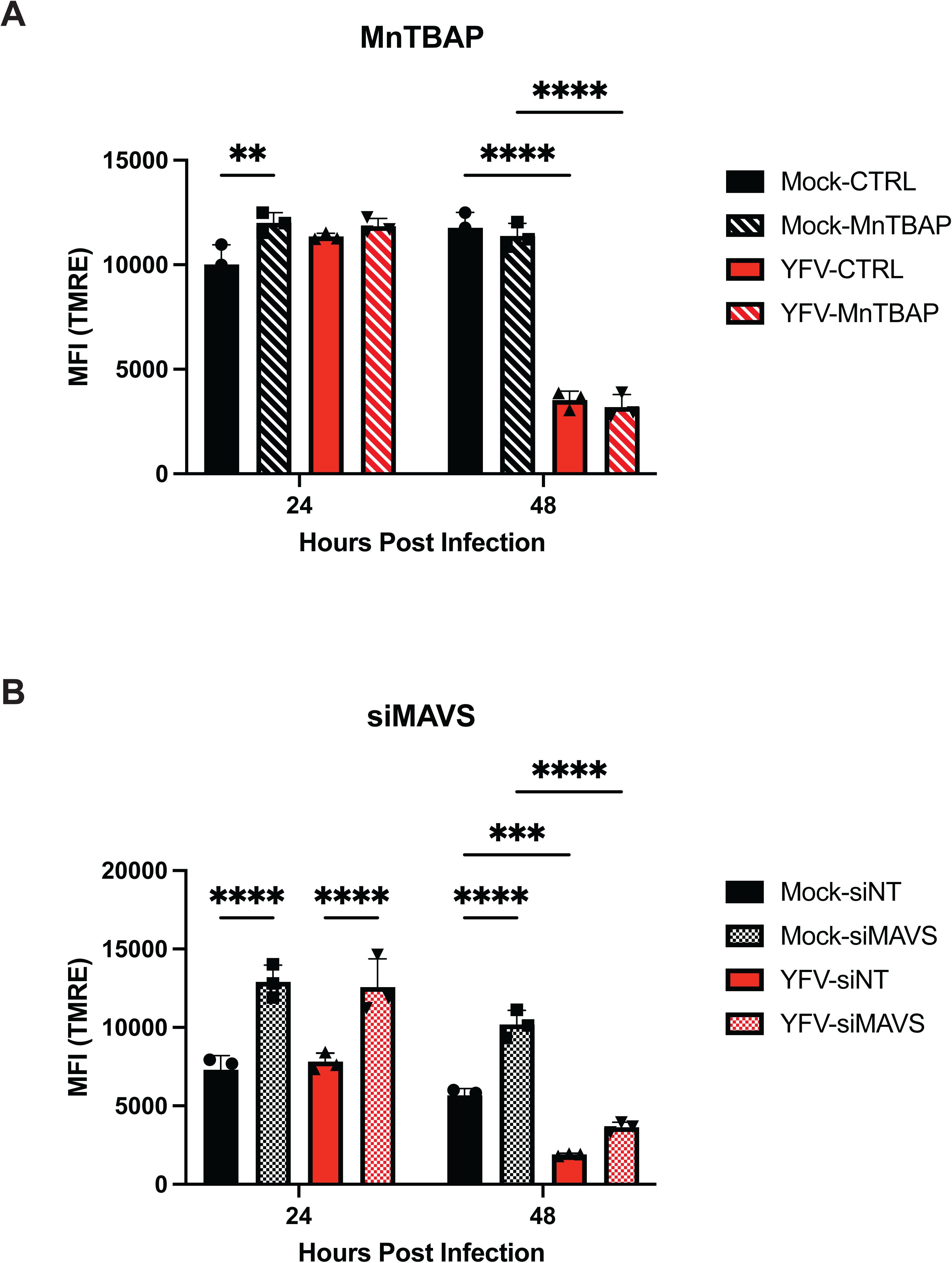
Loss of mitochondrial membrane potential is independent of type-I IFN signaling. Quantification of mitochondrial membrane potential (TMRE) in HepG2 cells infected with mock or YFV-17D (MOI 0.1) treated with **A** MnTBAP or **B** siMAVS to inhibit type-I IFN signaling. Values represent the mean ± SD (n=3). Statistical significance was assessed using two-way ANOVA followed by Tukey’s post hoc test for multiple groups. ** p<0.01, *** p<0.001, **** p<0.0001.

**Supplemental Figure 8:**
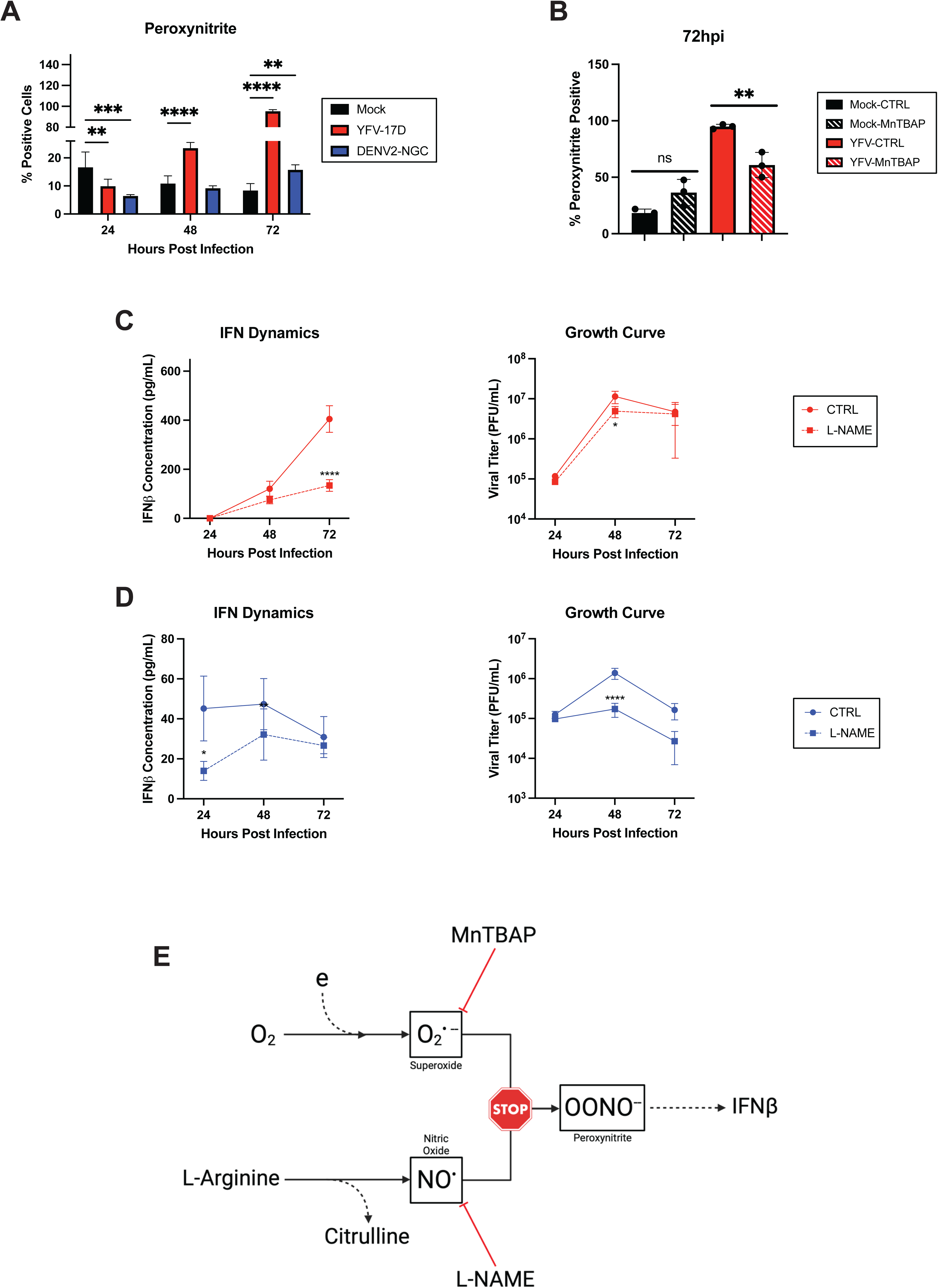
Peroxynitrite appears to be the main source of ROS in YFV-17D infection. **A** Quantification of peroxynitrite staining in mock, YFV-17D (MOI 0.1), and DENV2 (MOI 1) infected HepG2 cells at 24hpi, 48hpi, and 72hpi using flow cytometry. **B** Flow cytometric quantification of peroxynitrite staining in mock, YFV-17D, and DENV2 infected HepG2 cells treated with MnTBAP or a vehicle control for 48 hours. **C and D** Growth curve quantifying the viral titer and ELISA quantification of IFNβ secreted by HepG2 cells infected with **C.** YFV-17D or **D** DENV2 and treated with L-NAME [5mM] at 0hpi. **E** Schematic showing the proposed mechanism by which both MnTBAP and L-NAME inhibit peroxynititre formation therby blocking type-I IFN production. Values represent the mean ± SD (n=3). Statistical significance was assessed using two-way ANOVA followed by Dunnett’s multiple comparisons test (A), ordinary one-way ANOVA followed by Tukey’s multiple comparisons test (B), or two-way ANOVA followed by Sidak’s multiple comparisons test (C, D). **** p<0.0001.

**Supplemental Figure 9:**
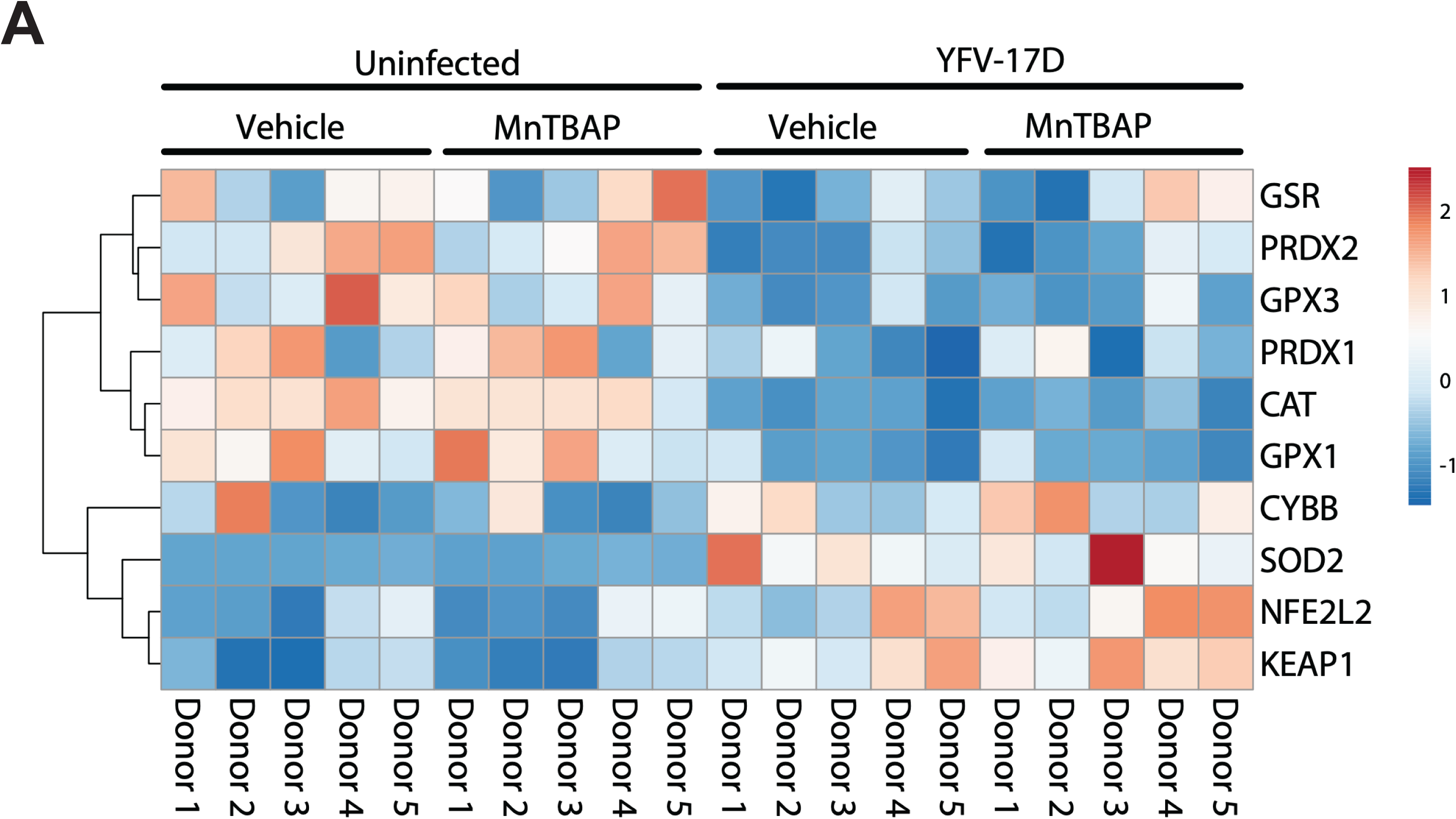
MnTBAP treatment ameliorates effects of YFV-17D infection. **A** Heatmap comparing the expression of oxidative stress genes in mock and YFV-17D (MOI 0.1) infected human DCs treated with and without MnTBAP.

---

**Supplemental Table S1.**

| Pathway | Enrichment Score | P-Value | Adjusted P-Value | Leading Edge Genes |
| --- | --- | --- | --- | --- |
| TRAF6 mediated IRF7 activation | -0.77 | 0.002 | 0.06 | IFNA10;IFNA5;IFNA8;IFNA21;IFNA2; IRF7;IFNA7;IFNA14;SIKE1;IFNA1;TANK; TRAF2;TBK1;IFIH1;TRIM25;IFNB1;IKBKE; RNF135 |
| Interferon alpha/beta signaling | -0.766 | 0.002 | 0.06 | IFIT2;IFIT1;USP18;RSAD2;ISG20;IFNA10; ISG15;IFIT3;SOCS1;IFI27;OAS1;IFNA5; IRF2;IFNA8;IFITM1;IFITM2;IFNA21;IRF9; IFI35;IFNA2;OASL;IRF7;IFNA7;IFIT5; IFNA14;IFITM3;STAT2;STAT1;BST2;IRF8; IRF1;IFNA1;GBP2;MX1;IP6K2;MX2; HLA-B;PSMB8;OAS2 |
| Regulation of IFNA/IFNB signaling | -0.722 | 0.002 | 0.06 | USP18;IFNA10;SOCS1;IFNA5;IFNA8; IFNA21;IFNA2;IFNA7;IFNA14;STAT2; IFNA1 |
| Interferon gamma signaling | -0.652 | 0.002 | 0.06 | GBP5;MT2A;GBP4;GBP1;SOCS1;OAS1; IRF2;FCGR1A;TRIM6;TRIM31;IRF9; TRIM21;TRIM38;TRIM22;OASL;SP100; IRF7;TRIM5;IRF8;IRF1;GBP3;VCAM1; GBP2; TRIM35;IFI30;TRIM26; HLA-B;OAS2 |
| Interferon signaling | -0.586 | 0.002 | 0.06 | IFIT2;GBP5;IFIT1;USP18;MT2A;GBP4; RSAD2;ISG20;GBP1;IFNA10;ISG15;IFIT3; SOCS1;IFI27;KPNA5;OAS1;IFNA5;IRF2; IFNA8;FCGR1A;TRIM6;TRIM31;IFITM1; IFITM2;IFNA21;IRF9;TRIM21;DDX58; IFI35;IFNA2;TRIM38;UBE2L6;TRIM22; OASL;SP100;IRF7;IFNA7;IFIT5;IFNA14; IFITM3;TRIM5;STAT2;EIF4E3;STAT1;BST2; IRF8;IRF1;IFNA1;GBP3;VCAM1;GBP2; TRIM35;MX1;IFI30;TRIM26;IP6K2;MX2; NUP160;HLA-B;PSMB8;UBA7;OAS2 |
| DDX58/IFIH1 mediated induction interferon alpha/beta | -0.574 | 0.002 | 0.06 | IFNA10;ISG15;IFNA5;IFNA8;IFNA21; DDX58;IFNA2;UBE2L6;IRF7;IFNA7;FADD; IFNA14;DHX58;SIKE1;IFNA1;TANK; ATG12;TRAF2;UBA7;TBK1;IFIH1;TRIM25; HERC5;UBE2D3;IFNB1;IKBKE;RNF135; UBE2D1;RIPK1 |

## References

1. Pierson, T.C. and M.S. Diamond, The continued threat of emerging flaviviruses. Nat Microbiol, 2020. 5(6): p. 796–812.

2. Koprowski, H. and M.B.A. Oldstone, Microbe hunters, then and now. 1996, Bloomington, Ill.: Medi-Ed Press. 456 p.

3. White, C.R., Yellow fever; history of the disease in the eighteenth and nineteenth century. J Kans Med Soc, 1959. 60(8): p. 298–302 passim.

4. Patterson, K.D., *Yellow fever epidemics and mortality in the United States*, *1693-1905*. Soc Sci Med, 1992. 34(8): p. 855–65.

5. Clements, A.N. and R.E. Harbach, History of the discovery of the mode of transmission of yellow fever virus. J Vector Ecol, 2017. 42(2): p. 208–222.

6. Theiler, M. and H.H. Smith, The effect of prolonged cultivation in vitro upon the pathogenicity of yellow fever virus. Journal of Experimental Medicine, 1937. 65(6): p. 767–786.

7. Theiler, M. and H.H. Smith, The use of yellow fever virus modified by in vitro cultivation for human immunization. Journal of Experimental Medicine, 1937. 65(6): p. 787–800.

8. Monath, T.P. and P.F. Vasconcelos, Yellow fever. J Clin Virol, 2015. 64: p. 160–73.

9. Pulendran, B., Learning immunology from the yellow fever vaccine: innate immunity to systems vaccinology. Nat Rev Immunol, 2009. 9(10): p. 741–7.

10. Barrett, A.D. and D.E. Teuwen, Yellow fever vaccine - how does it work and why do rare cases of serious adverse events take place? Curr Opin Immunol, 2009. 21(3): p. 308–13.

11. Gaucher, D., et al., Yellow fever vaccine induces integrated multilineage and polyfunctional immune responses. J Exp Med, 2008. 205(13): p. 3119–31.

12. Akondy, R.S., et al., Initial viral load determines the magnitude of the human CD8 T cell response to yellow fever vaccination. Proc Natl Acad Sci U S A, 2015. 112(10): p. 3050–5.

13. Pulendran, B., Variegation of the immune response with dendritic cells and pathogen recognition receptors. J Immunol, 2005. 174(5): p. 2457–65.

14. Steinman, R.M. and J. Banchereau, Taking dendritic cells into medicine. Nature, 2007. 449(7161): p. 419–26.

15. Takeuchi, O. and S. Akira, Innate immunity to virus infection. Immunol Rev, 2009. 227(1): p. 75–86.

16. Douam, F., et al., Type III Interferon-Mediated Signaling Is Critical for Controlling Live Attenuated Yellow Fever Virus Infection In Vivo. mBio, 2017. 8(4).

17. Lam, L.K.M., et al., Gamma-interferon exerts a critical early restriction on replication and dissemination of yellow fever virus vaccine strain 17D-204. NPJ Vaccines, 2018. 3: p. 5.

18. Querec, T., et al., *Yellow fever vaccine YF-17D activates multiple dendritic cell subsets via TLR2*, *7*, *8, and 9 to stimulate polyvalent immunity*. J Exp Med, 2006. 203(2): p. 413–24.

19. Douam, F. and A. Ploss, Yellow Fever Virus: Knowledge Gaps Impeding the Fight Against an Old Foe. Trends Microbiol, 2018. 26(11): p. 913–928.

20. Bastard, P., et al., Auto-antibodies to type I IFNs can underlie adverse reactions to yellow fever live attenuated vaccine. J Exp Med, 2021. 218(4).

21. Hernandez, N., et al., Inherited IFNAR1 deficiency in otherwise healthy patients with adverse reaction to measles and yellow fever live vaccines. J Exp Med, 2019. 216(9): p. 2057–2070.

22. Suthar, M.S., et al., A systems biology approach reveals that tissue tropism to West Nile virus is regulated by antiviral genes and innate immune cellular processes. PLoS Pathog, 2013. 9(2): p. e1003168.

23. Rehwinkel, J. and M.U. Gack, RIG-I-like receptors: their regulation and roles in RNA sensing. Nat Rev Immunol, 2020. 20(9): p. 537–551.

24. Chatel-Chaix, L., et al., Dengue Virus Perturbs Mitochondrial Morphodynamics to Dampen Innate Immune Responses. Cell Host Microbe, 2016. 20(3): p. 342–356.

25. Aguirre, S., et al., Dengue virus NS2B protein targets cGAS for degradation and prevents mitochondrial DNA sensing during infection. Nat Microbiol, 2017. 2: p. 17037.

26. Aguirre, S., et al., DENV inhibits type I IFN production in infected cells by cleaving human STING. PLoS Pathog, 2012. 8(10): p. e1002934.

27. Yu, C.Y., et al., Dengue virus targets the adaptor protein MITA to subvert host innate immunity. PLoS Pathog, 2012. 8(6): p. e1002780.

28. Chan, K.R., et al., Metabolic perturbations and cellular stress underpin susceptibility to symptomatic live-attenuated yellow fever infection. Nat Med, 2019. 25(8): p. 1218–1224.

29. Querec, T.D., et al., Systems biology approach predicts immunogenicity of the yellow fever vaccine in humans. Nat Immunol, 2009. 10(1): p. 116–125.

30. Fernandez-Garcia, M.D., et al., Vaccine and Wild-Type Strains of Yellow Fever Virus Engage Distinct Entry Mechanisms and Differentially Stimulate Antiviral Immune Responses. mBio, 2016. 7(1): p. e01956–15.

31. Sun, B., et al., Dengue virus activates cGAS through the release of mitochondrial DNA. Sci Rep, 2017. 7(1): p. 3594.

32. Castanier, C., et al., Mitochondrial dynamics regulate the RIG-I-like receptor antiviral pathway. EMBO Rep, 2010. 11(2): p. 133–8.

33. Onoguchi, K., et al., Virus-infection or 5’ppp-RNA activates antiviral signal through redistribution of IPS-1 mediated by MFN1. PLoS Pathog, 2010. 6(7): p. e1001012.

34. Plitzko, B. and S. Loesgen, Measurement of Oxygen Consumption Rate (OCR) and Extracellular Acidification Rate (ECAR) in Culture Cells for Assessment of the Energy Metabolism. Bio Protoc, 2018. 8(10): p. e2850.

35. Mills, E.L., B. Kelly, and L.A.J. O’Neill, Mitochondria are the powerhouses of immunity. Nat Immunol, 2017. 18(5): p. 488–498.

36. Muri, J. and M. Kopf, Redox regulation of immunometabolism. Nat Rev Immunol, 2021. 21(6): p. 363–381.

37. Pacher, P., J.S. Beckman, and L. Liaudet, Nitric oxide and peroxynitrite in health and disease. Physiol Rev, 2007. 87(1): p. 315–424.

38. Hagan, T., et al., Transcriptional atlas of the human immune response to 13 vaccines reveals a common predictor of vaccine-induced antibody responses. Nat Immunol, 2022. 23(12): p. 1788–1798.

39. Kasturi, S.P., et al., Programming the magnitude and persistence of antibody responses with innate immunity. Nature, 2011. 470(7335): p. 543–7.

40. Li, S., et al., Metabolic Phenotypes of Response to Vaccination in Humans. Cell, 2017. 169(5): p. 862–877 e17.

41. Ravindran, R., et al., Vaccine activation of the nutrient sensor GCN2 in dendritic cells enhances antigen presentation. Science, 2014. 343(6168): p. 313–317.

42. Ponia, S.S., et al., Mitophagy antagonism by ZIKV reveals Ajuba as a regulator of PINK1 signaling, PKR-dependent inflammation, and viral invasion of tissues. Cell Rep, 2021. 37(4): p. 109888.

43. Sorouri, M., T. Chang, and D.C. Hancks, Mitochondria and Viral Infection: Advances and Emerging Battlefronts. mBio, 2022. 13(1): p. e0209621.

44. Khan, M., et al., Mitochondrial dynamics and viral infections: A close nexus. Biochim Biophys Acta, 2015. 1853(10 Pt B): p. 2822–33.

45. Tiku, V., M.W. Tan, and I. Dikic, Mitochondrial Functions in Infection and Immunity. Trends Cell Biol, 2020. 30(4): p. 263–275.

46. Koshiba, T., et al., Mitochondrial membrane potential is required for MAVS-mediated antiviral signaling. Sci Signal, 2011. 4(158): p. ra7.

47. Frierson, J.G., The yellow fever vaccine: a history. Yale J Biol Med, 2010. 83(2): p. 77–85.

48. Tur, J., et al., Mitofusin 2 in Macrophages Links Mitochondrial ROS Production, Cytokine Release, Phagocytosis, Autophagy, and Bactericidal Activity. Cell Rep, 2020. 32(8): p. 108079.

49. Chen, S., et al., TBK1-Mediated DRP1 Targeting Confers Nucleic Acid Sensing to Reprogram Mitochondrial Dynamics and Physiology. Mol Cell, 2020. 80(5): p. 810–827 e7.

50. O’Carroll, S.M., F.D.R. Henkel, and L.A.J. O’Neill, Metabolic regulation of type I interferon production. Immunol Rev, 2024. 323(1): p. 276–287.

51. Everts, B., et al., TLR-driven early glycolytic reprogramming via the kinases TBK1-IKKvarepsilon supports the anabolic demands of dendritic cell activation. Nat Immunol, 2014. 15(4): p. 323–32.

52. O’Neill, L.A. and E.J. Pearce, Immunometabolism governs dendritic cell and macrophage function. J Exp Med, 2016. 213(1): p. 15–23.

53. Olagnier, D., et al., Cellular oxidative stress response controls the antiviral and apoptotic programs in dengue virus-infected dendritic cells. PLoS Pathog, 2014. 10(12): p. e1004566.

54. Wang, R., et al., Influenza M2 protein regulates MAVS-mediated signaling pathway through interacting with MAVS and increasing ROS production. Autophagy, 2019. 15(7): p. 1163–1181.

55. Buskiewicz, I.A., et al., Reactive oxygen species induce virus-independent MAVS oligomerization in systemic lupus erythematosus. Sci Signal, 2016. 9(456): p. ra115.

56. Nobre, L., et al., Modulation of Innate Immune Signalling by Lipid-Mediated MAVS Transmembrane Domain Oligomerization. PLoS One, 2015. 10(8): p. e0136883.

57. Thannickal, V.J. and B.L. Fanburg, Reactive oxygen species in cell signaling. Am J Physiol Lung Cell Mol Physiol, 2000. 279(6): p. L1005–28.

58. Zhang, H., et al., Transmembrane nitration of hydrophobic tyrosyl peptides. Localization, characterization, mechanism of nitration, and biological implications. J Biol Chem, 2003. 278(11): p. 8969–78.

59. Greenacre, S.A. and H. Ischiropoulos, Tyrosine nitration: localisation, quantification, consequences for protein function and signal transduction. Free Radic Res, 2001. 34(6): p. 541–81.

60. Zhou, H.L., et al., An enzyme that selectively S-nitrosylates proteins to regulate insulin signaling. Cell, 2023. 186(26): p. 5812–5825 e21.

61. Gauba, V., et al., Loss of CD4 T-cell-dependent tolerance to proteins with modified amino acids. Proc Natl Acad Sci U S A, 2011. 108(31): p. 12821–6.

62. Turko, I.V., et al., Protein tyrosine nitration in the mitochondria from diabetic mouse heart. Implications to dysfunctional mitochondria in diabetes. J Biol Chem, 2003. 278(36): p. 33972–7.

63. Hahn, C.S., et al., Comparison of the virulent Asibi strain of yellow fever virus with the 17D vaccine strain derived from it. Proc Natl Acad Sci U S A, 1987. 84(7): p. 2019–23.

64. Lee, E. and M. Lobigs, E protein domain III determinants of yellow fever virus 17D vaccine strain enhance binding to glycosaminoglycans, impede virus spread, and attenuate virulence. J Virol, 2008. 82(12): p. 6024–33.

65. McCloskey, D., et al., A pH and solvent optimized reverse-phase ion-paring-LC-MS/MS method that leverages multiple scan-types for targeted absolute quantification of intracellular metabolites. Metabolomics, 2015. 11(5): p. 1338–1350.

66. Groveman, B.R., et al., A PrP EGFR signaling axis controls neural stem cell senescence through modulating cellular energy pathways. J Biol Chem, 2023. 299(11): p. 105319.

67. Gao, Z., et al., Development of antibody-based assays for high throughput discovery and mechanistic study of antiviral agents against yellow fever virus. Antiviral Res, 2020. 182: p. 104907.

68. Sanchez-Velazquez, R., et al., Generation of a reporter yellow fever virus for high throughput antiviral assays. Antiviral Res, 2020. 183: p. 104939.

69. Cheung, A.M., et al., Characterization of Live-Attenuated Powassan Virus Vaccine Candidates Identifies an Efficacious Prime-Boost Strategy for Mitigating Powassan Virus Disease in a Murine Model. Vaccines (Basel), 2023. 11(3).

70. Shannon, J.G., D. Howe, and R.A. Heinzen, Virulent Coxiella burnetii does not activate human dendritic cells: role of lipopolysaccharide as a shielding molecule. Proc Natl Acad Sci U S A, 2005. 102(24): p. 8722–7.

71. Dobin, A., et al., STAR: ultrafast universal RNA-seq aligner. Bioinformatics, 2013. 29(1): p. 15–21.

72. Li, B. and C.N. Dewey, RSEM: accurate transcript quantification from RNA-Seq data with or without a reference genome. BMC Bioinformatics, 2011. 12: p. 323.

73. Ritchie, M.E., et al., *limma powers differential expression analyses for RNA-sequencing and microarray studies*. Nucleic Acids Res, 2015. 43(7): p. e47.

74. Subramanian, A., et al., Gene set enrichment analysis: a knowledge-based approach for interpreting genome-wide expression profiles. Proc Natl Acad Sci U S A, 2005. 102(43): p. 15545–50.

75. Milacic, M., et al., The Reactome Pathway Knowledgebase 2024. Nucleic Acids Res, 2024. 52(D1): p. D672–D678.

76. Rath, S., et al., MitoCarta3.0: an updated mitochondrial proteome now with sub-organelle localization and pathway annotations. Nucleic Acids Res, 2021. 49(D1): p. D1541–D1547.

